# The accessible chromatin landscape of the hippocampus at single-cell resolution

**DOI:** 10.1101/407668

**Authors:** John R. Sinnamon, Kristof A. Torkenczy, Michael W. Linhoff, Sarah Vitak, Hannah A. Pliner, Cole Trapnell, Frank J. Steemers, Gail Mandel, Andrew C. Adey

**Affiliations:** The Vollum Institute, Oregon Health & Science University, Portland, OR, USA; Oregon Health & Science University, Department of Molecular and Medical Genetics, Portland, OR, USA; University of Washington, Department of Genome Sciences, Seattle, WA, USA; Illumina, Inc., San Diego, CA, USA; Knight Cardiovascular Institute, Oregon Health & Science University, Portland, OR, USA; Knight Cancer Institute, Oregon Health & Science University, Portland, OR, USA

## Abstract

Here we present a comprehensive map of the accessible chromatin landscape of the mouse hippocampus at single-cell resolution. Substantial advances of this work include the optimization of single-cell combinatorial indexing assay for transposase accessible chromatin (sci-ATAC-seq), a software suite, *scitools*, for the rapid processing and visualization of single-cell combinatorial indexing datasets, and a valuable resource of hippocampal regulatory networks at single-cell resolution. We utilized sci-ATAC-seq to produce 2,346 high-quality single-cell chromatin accessibility maps with a mean unique read count per cell of 29,201 from both fresh and frozen hippocampi, observing little difference in accessibility patterns between the preparations. Using this dataset, we identified eight distinct major clusters of cells representing both neuronal and non-neuronal cell types and characterized the driving regulatory factors and differentially accessible loci that define each cluster. We then applied a recently described co-accessibility framework, *Cicero*, which identified 146,818 links between promoters and putative distal regulatory DNA. Identified co-accessibility networks showed cell-type specificity, shedding light on key dynamic loci that reconfigure to specify hippocampal cell lineages. Lastly, we carried out an additional sci-ATAC-seq preparation from cultured hippocampal neurons (899 high-quality cells, 43,532 mean unique reads) that revealed substantial alterations in their epigenetic landscape compared to nuclei from hippocampal tissue. This dataset and accompanying analysis tools provide a new resource that can guide subsequent studies of the hippocampus.

## Introduction

A major goal in the life sciences is to map cell types and identify the respective genomic properties of each of the cell types in complex tissues. Traditional strategies that utilize intact tissue are limited to averaging of the constituent cell profiles. An alternative approach is to target specific cell populations via antibody-based or reporter gene labeling; however, these strategies are limited by the small number of markers that can be utilized during cell isolation, and the markers usually label more than one cell population within the tissue. Additionally, the selected markers must be known in advance. All of these limitations are exacerbated when the tissue of interest has a high degree of cellular complexity, as in the brain. To overcome these limitations, there has been a burst in development of unbiased single-cell genomics assays, leveraging the concept that each single cell can only occupy a single position in the landscape of cell types (Trapnell 2015).

This push into the single-cell space has largely centered on the use of single-cell transcriptional profiling, where high cell count strategies that produce enough information per cell to deconvolve cell types are typically deployed. While profiling the RNA complement has produced valuable information (Zeisel et al. 2018; Saunders et al. 2018), the ability to profile chromatin status, *i.e.* active versus inactive, has lagged behind, leaving open the question as to what extent accessible chromatin profiles are linked to cell specificity, particularly with respect to distal enhancer elements (Corces et al. 2016).

Recently, progress has been made to ascertain chromatin accessibility profiles in single cells using ATAC-seq (Assay for Transposase-Accessible Chromatin) technologies. These strategies have been applied to myogenesis (Pliner et al. 2018), hematopoietic differentiation (Buenrostro et al. 2018), fly embryonic development (Cusanovich et al. 2018b), the mouse (Preissl et al. 2018) and human cortex (Lake et al. 2017), and most recently an atlas of multiple tissues in the mouse, though notably lacking the hippocampus (Cusanovich et al. 2018a). The core concept behind the methods utilized in several of these studies is a combinatorial indexing schema whereby library molecules are barcoded twice, once at the transposase stage and then again at the PCR stage. Initially developed for the acquisition of long-range sequence information, *i.e.* linked-reads for purposes resolving the haplotypes of genomic variants or for *de novo* genome assembly (Amini et al. 2014; Adey et al. 2014), this barcoding technology is the key component for single-cell combinatorial indexing ATAC-seq, or sci-ATAC-seq (Cusanovich et al. 2015). This platform has also been extended to profile other properties including transcription, genome sequencing, chromatin folding, and DNA methylation (Cao et al. 2017; Ramani et al. 2017; Vitak et al. 2017; Mulqueen et al. 2018; Yin et al. 2018). In this work, we optimized the sci-ATAC-seq assay for analysis of fresh and frozen hippocampal tissue samples to produce single-cell chromatin accessibility profiles in high throughput, with greater information content – as measured by unique reads per cell – when compared to previous high-throughput methods, and comparable to microfluidics-based technologies (Buenrostro et al. 2015). These improvements will also facilitate the use of this technology platform on frozen samples, enabling the assessment of banked tissue isolates.

The hippocampus is critical to the formation and retrieval of episodic and spatial memory (Zola-Morgan et al. 1986; Smith and Milner 1981; O’Keefe and Dostrovsky 1971; Scoville and Milner 1957). Historically, cell types within the hippocampus have been broadly classified by their morphology (Ramon y Cajal 1911; Lorente de No 1934) and electrophysiological properties (Spencer and Kandel 1961b, 1961a; Kandel and Spencer 1961; Kandel et al. 1961). More recently, transcriptional profiling has demonstrated cell types can be identified by their transcriptional profiles (Lein et al. 2004; Cembrowski et al. 2016), and single-cell transcriptomics has also revealed potential subclasses within previously defined cell types (Habib et al. 2017; Zeisel et al. 2015). The defined classes of cells within the hippocampus and the existing single cell transcriptome data allowed us to refine our sci-ATAC-seq method and provide the first single-cell epigenomics profile of the murine hippocampus.

## Results

### Single-cell chromatin accessibility profiles from mouse hippocampus

We implemented an improved sci-ATAC-seq protocol on two fresh and two frozen mouse hippocampi to map the accessible chromatin landscape (Methods). Each sample was freshly isolated from an adult (P60) wild type mouse (C57-Bl6) and either processed immediately or flash frozen using liquid nitrogen. Nuclei were isolated and carried through the sci-ATAC-seq protocol (Methods, Fig 1A). Briefly, nuclei were distributed by Fluorescence Assisted Nuclei Sorting (FANS) using DAPI as a stain to select for intact, single nuclei. We deposited 2,500 nuclei into each well of a 96-well plate containing transposition reaction buffer followed by the addition of the transposase enzyme loaded with DNA sequencing adaptors containing barcodes unique to each well of the plate. The transposome complexes are able to freely enter into the nucleus and insert into regions of open chromatin, as in standard ATAC-seq, without disrupting nuclear scaffold integrity. We then pooled all 96 wells and performed FANS again to distribute 22 nuclei into each well of a new 96-well plate for subsequent PCR with primers containing additional sequence barcodes corresponding to each well of the PCR plate. Each resulting molecule produces a sequence read with two barcode sets – one for the transposase stage of indexing, and the second for the PCR stage – which enables the unique cell identity of the sequence read.

**Figure 1.**
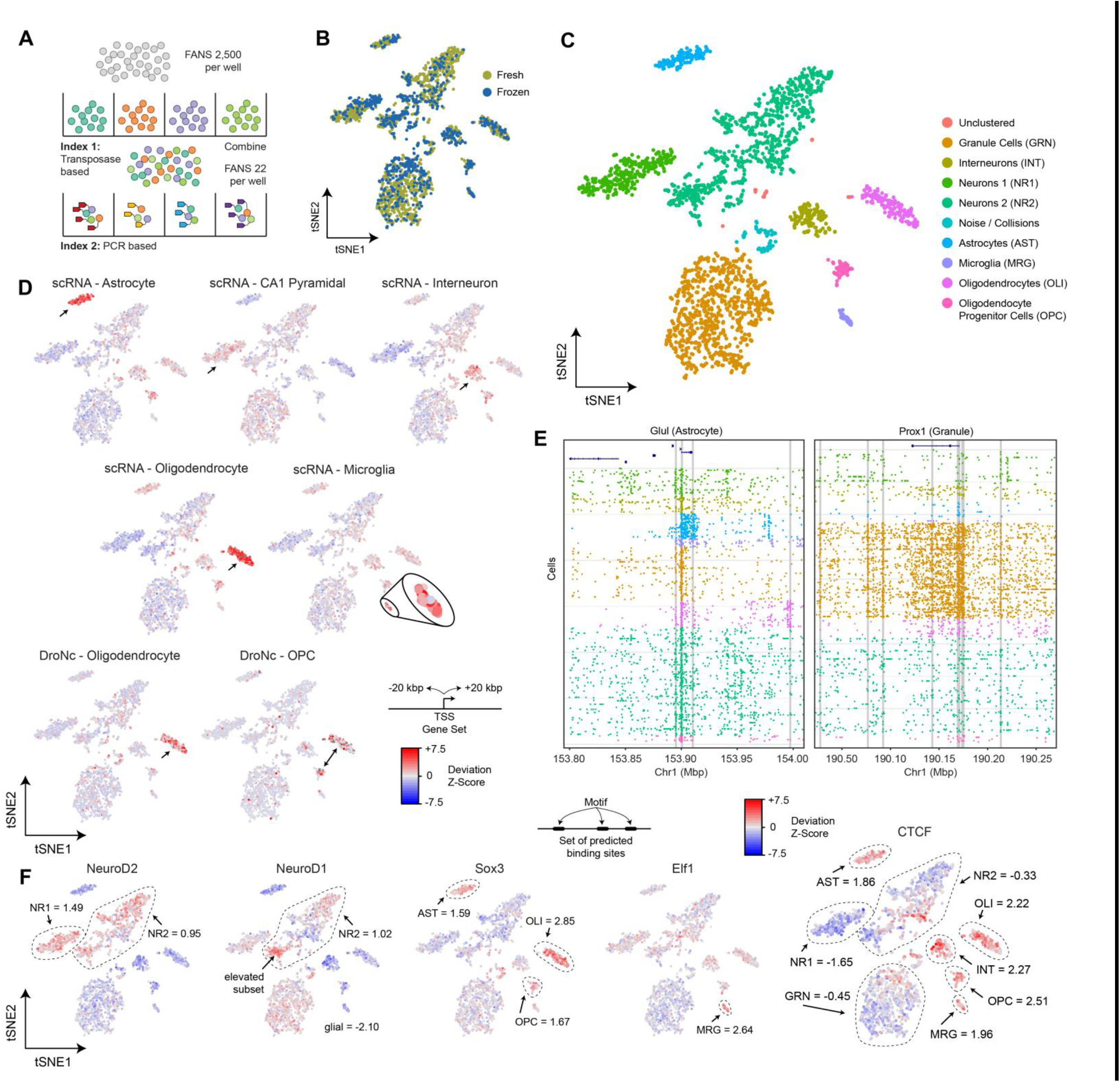
sci-ATAC-seq of the murine hippocampus. (A) sci-ATAC-seq workflow. Two indexes are incorporated into library molecules for each cell enabling single-cell discrimination. (B) LSI-tSNE projection of single cells colored by tissue preparation method. Little variation in tSNE space is observed between fresh or frozen starting material. (C) LSI-tSNE projection of cells colored by assigned cluster and cell type. Enrichment of accessibility of proximal regulatory elements for marker genes as identified by single-cell RNA-seq (Zeisel et al. 2015) and DroNc-seq (Habib et al. 2017) for each cell. The microglial population is enlarged for visibility. Black arrows indicate the cell cluster associated with the marker gene set. (E) sci-ATAC-seq read plots at *Glul* (astrocyte marker gene) and *Prox1* (dentate granule cell marker gene). (F) ChromVAR global motif deviation z-scores for each cell for select motifs. Dashed lines and values correspond to mean values of cell populations.

In total, we produced 2,346 single cells passing quality control (≥1,000 and ≥25% unique reads present in called peaks) evenly represented across replicates (2 frozen, 2 fresh; Supplementary Table 1). Cells had a mean unique aligned read count of 29,201, which is notably higher than other high throughput single-cell ATAC-seq workflows to date (generally ~10,000). We observed a strong correlation in ATAC signal between the aggregate profiles of the four replicates (Pearson R > 0.99), indicating high reproducibility across preparations for both fresh and frozen tissue. Chromatin accessibility peaks were identified by the aggregation of all cells to produce an ensemble dataset containing all called peaks, resulting in a preliminary set of 93,994 high-confidence peaks, with a mean of 36.4% of reads from each cell falling within these regions.

We constructed a read count matrix of our ensemble peaks and single cells from all conditions (Supplementary Data – InVivo.counts.matrix) by tallying the number of reads for each cell at each peak. We next utilized *scitools* to perform Latent Semantic Indexing (LSI), as previously described (Cusanovich et al. 2015, 2018b), with the exclusion of cells with reads at fewer than 1,000 sites and of sites with fewer than 50 cells exhibiting signal. The LSI matrix was projected into two-dimensional space using t-distributed Stochastic Neighbor Embedding (tSNE) for visualization, which revealed distinct domains occupied by clusters of cells. We then used a density based method (Ester et al. 1996) to identify nine major clusters (Fig. 1C), one of which was identified to be likely barcode collisions and removed from further analysis (Methods). A comparison of the proportion of cells assigned to each cluster with respect to fresh or frozen samples did not yield a significant difference (X^2^ = 9.85, *p*-value = 0.20; Fig. 1B, Supplementary Table 1), though increased proportions of interneurons and microglia were observed in the frozen preparation. This encouraging finding underscores the robustness of our sci-ATAC-seq workflow and will enable experiments for which only frozen samples are available.

To assign each of our identified clusters to a cell type, we took advantage of published single-cell RNA-seq data that produced sets of marker genes associated with cell types identified at the transcriptional level (Zeisel et al. 2015; Habib et al. 2017). Assuming that regulatory sequences proximal to these genes are likely be enriched for elements that drive cell-specific expression, cells should exhibit increased levels of chromatin accessibility for the set of peaks associated with the marker genes of their respective cell type. For each set of cell-type-specific genes, we identified peaks 20 kilobasepairs (kbp) in either direction from the transcriptional start site. These peak sets were used to calculate the enrichment for accessible chromatin for each cell within these regions to produce a deviation z-score, similar to previously described methods (Buenrostro et al. 2015; Schep et al. 2017). We then visualized these scores on our tSNE projections, which enabled us to clearly identify a number of neuronal and non-neuronal cell types, including astrocytes (AST), two groups of pyramidal neurons (NR1 and NR2), interneurons (INT), oligodendrocytes (OLI), microglia (MRG), and oligodendrocyte progenitor cells (OPCs) (Fig. 1D). However, this enrichment-based strategy is heavily reliant on a robust set of marker genes that are presumed to lack expression in all other cell types. To complement this strategy we also turned to marker genes described previously in the literature that were not present in available single-cell RNA-seq datasets and assessed the chromatin accessibility at elements proximal to these genes (Fig. 1E, Supplementary Fig. 1). For example, the *Glul* gene, an established marker for astrocytes (Martinez-Hernandez et al. 1977; Fages et al. 1988) showed accessibility only in the population of cells we identified as astrocytes (Fig. 1E, left). *Prox1,* previously shown to be enriched in the dentate gyrus (Lein et al. 2004), is accessible predominantly in the dentate granule cell population (GRN, Fig. 1E, right). Markers for particular cell types were also consistent with *in situ* hybridization data from the Allen Brain Institute (Supplementary Fig. 1) and RNA-seq data from sorted cells (Cembrowski et al. 2016; Zhang et al. 2014).

Based on our cell type assignments, the number of cells in each population reflects the proportions seen within the intact hippocampus. For example, the two major cell types, by cell number according to stereological estimates, are the excitatory pyramidal cells and dentate granule cells, with each population consisting of approximately 400,000 cells per mouse hippocampus (Abusaad et al. 1999). Using cell type specific markers, it has been estimated that within subregions of the mouse hippocampus, there are 10 to 14 fold fewer astrocytes and 42 to 74 fold fewer microglia compared to neurons (Kimoto et al. 2009), which is consistent with our observation of 14 fold and 41 fold fewer astrocytes and microglia, respectively (Supplementary Table 1).

### Global DNA binding motif accessibility

Chromatin accessibility maps largely represent the active (open) enhancer and promoter landscape. These regulatory elements typically harbor sets of motifs recognized by DNA binding proteins that in turn recruit histone modifiers, e.g. acetylases, deacetylases, methylases, to ultimately dictate the transcriptional status of genes. To assess the global activity of DNA binding proteins we utilized the recently-described software tool, *ChromVAR* (Schep et al. 2017), which aggregates the chromatin accessibility signal genome-wide at sites harboring a given motif, followed by the calculation of a deviation z-score for each cell. This score represents the putative activity level of the DNA binding protein that corresponds to the assessed motif, which we then visualized on our tSNE projections (Fig. 1F). Although motif accessibility for a particular DNA binding protein can be confounded by similar motifs or motifs that co-occur at the same regulatory loci, the ability to assess any given motif for accessibility provides a powerful tool to identify targets for subsequent analysis.

In line with expectations, our cell type clusters showed enrichment for accessibility at DNA binding motifs concordant with the identified cell type (Fig. 1F, Supplementary Fig. 2). The analysis included the assessment of global accessibility for neuron-specific factors such NeuroD2, which associates with active chromatin marks (e.g. H3K27ac) in cortical tissue (Guner et al. 2017) and exhibited greater accessibility in the two pyramidal cell clusters (mean z-score (µ_z_) = 1.49 and 0.95 for NR1 and NR2 respectively, all other cell types µ_z_ ≤ −0.74). We also observed increased accessibility of NeuroD1, also associated with active chromatin (Pataskar et al. 2016), in a portion of one of the pyramidal neuron clusters (NR2, µ_z_ = 1.02) with less accessibility across glial populations (µ_z_ ≤ −2.10). While many studies have identified a role for Sox3 during neural differentiation, consistent with a previous expression study (Cheah and Thomas 2015), we observed increased Sox3 accessibility in astrocyte (µ_z_ = 1.59), oligodendrocyte (µ_z_ = 2.85), and OPC populations (µ_z_ = 1.67), suggesting a glial role for this transcription factor in adulthood. Elf-1, an ETS family member associated with activating interferon response in the hematopoietic lineage (Larsen et al. 2015), exhibited elevated accessibility in the microglial population (µ_z_ = 2.64), which also respond to interferon in the brain (e.g. (Goldmann et al. 2015)). Of particular interest was the strong enrichment for CTCF motif accessibility in glial cell populations (AST µ_z_ = 1.86, OLI µ_z_ = 2.22, OPC µ_z_ = 2.51, MRG µ_z_ = 1.96) and interneurons (µ = 2.27) when compared to granule cells (µ_z_ = −0.45) or pyramidal neurons (NR1 µ_z_ = −1.65, NR2 µ_z_ = −0.33), an observation that was reinforced by our subsequent differential accessibility analysis described below.

### Differential Accessibility by Cell Type

We next sought to show that accessible regions could be identified according to cell type. To provide sufficient signal, we aggregated cells within clusters in their local neighborhoods as has been described previously (Cusanovich et al. 2018b) and then carried out a differential accessibility analysis for each cluster compared to the rest of the cells (Methods, Fig. 2A). The power to detect cell-type-specific accessibility loci was tied largely to the number of cells within each population. However, each group produced a substantial number of passing hits (*q*-value ≤ 0.01, Log2 fold-change ≥ 1) ranging from 894 (OPCs) to 7,796 (granule cells). When assessing ATAC-seq signal for aggregated cells within each cell type at these sites, signal specificity was greatest for the two pyramidal neuron clusters with increased off-target signal for other non-neuronal populations, which had less signal (Fig. 2B, left, Supplementary Fig. 3-5).

**Figure 2.**
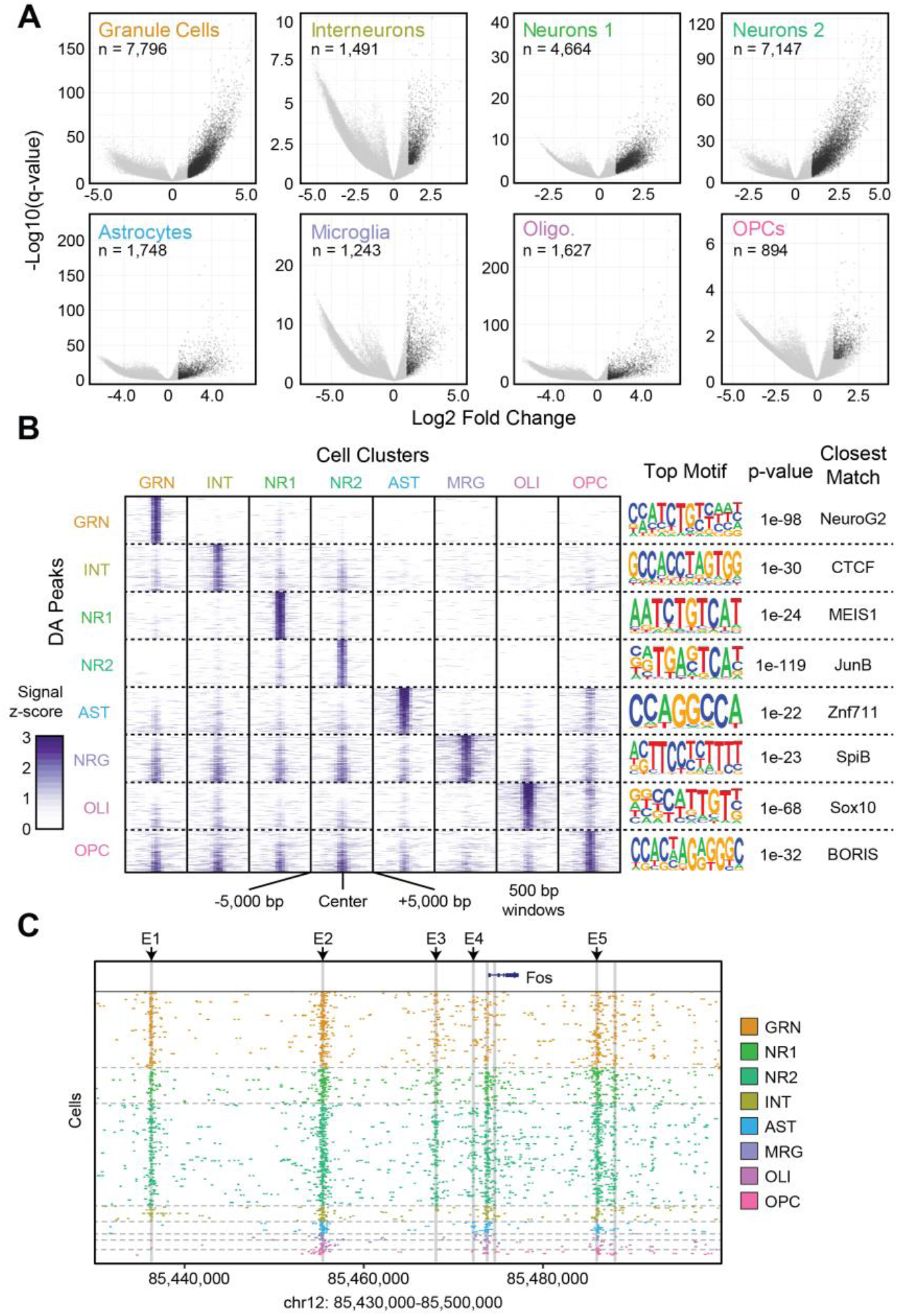
Differential accessibility analysis between cell types. (A) Volcano plots –log_10_(*q*-value) (y-axis) versus log2 accessibility signal fold change (x-axis) showing all peaks. Each comparison is for the indicated cell population versus all other cell types. Significant peaks (number indicated, *q*-value ≤ 0.01, log2 fold change ≥ 1) are in black. (B) ATAC-seq signal plots for the top differential accessible peaks for each cell type. The most significantly enriched motif for each set is shown on the right along with the corresponding *p*-value and closest matching known motif. (C) *c-Fos* locus with enhancers E1-5 highlighted to show cell-type-specific utilization.

We performed a motif enrichment analysis to identify DNA binding proteins that may bind within the differentially accessible regions (Fig. 2B, right). In contrast to the prior, global accessibility analysis, where all accessible loci were utilized to detect increased signal at sites harboring a given motif in each cell; here, we are detecting enrichment of motifs in the specific subsets of loci that were determined to be differentially accessible. This strategy revealed enrichment for binding by the Sox10 transcription factor in oligodendrocytes (Claus Stolt et al. 2002) and by NeuroG2 in the dentate granule cells (Roybon et al. 2009). Concordant with our global analysis, CTCF was enriched in the interneuron population. This is unexpected because CTCF is considered a general factor for regulation of chromatin architecture. To confirm the presence of CTCF binding in regions of differential accessibility in interneurons, we used publicly available CTCF Chromatin Immunoprecipitation sequencing (ChIP-seq) data from adult mouse hippocampus (Sams et al. 2016). One of the top regions of differential accessibility in interneurons was in an intron in the gene encoding actin filament associated protein 1 (*Afap1*, Supplementary Fig. 6). The ChIP data revealed CTCF binding within the same intron flanking the accessible region. While there have been no studies focused on the function of CTCF specifically in interneurons, there is evidence CTCF may have a particular importance in this cell type. CTCF binding motifs were enriched in the accessible chromatin of affinity purified parvalbumin positive cortical interneurons but not in VIP positive interneurons or excitatory neurons (Mo et al. 2015) and in mice expressing one CTCF allele only inhibitory neurons exhibit memory impairment (Kim et al. 2018). The potential selective importance of CTCF in interneurons warrants further study.

To further determine the utility of our method in assigning regulatory elements to cell types, we tested whether we could parse enhancers that had been identified in the literature as inducers of target genes in response to neuronal activity. We focused on the *c-Fos* gene that has been studied previously as a general reporter of neuronal activity throughout the brain. Specifically, five enhancers (E1-E5) have been characterized (Kim et al. 2010) for both regulation during neuronal activity and type of stimulation (Joo et al. 2015). Surprisingly, when we examined ATAC-seq signals at the five enhancers across cell types in hippocampus, we identified cell type specific patterns of accessibility. Notably, enhancers E1 and E3 were accessible only in neurons, while E2 and E5 were accessible in all cell types (Fig. 2C). Further, enhancer E4 was accessible in group 2 but not group 1 pyramidal neurons and was also accessible in a small portion of dentate granule cells. Our findings suggest cell type specificity in stimuli responsiveness within the hippocampus, even between pyramidal cell subpopulations, opening the door to new studies of the basis of these signaling differences and demonstrating the utility of single-cell epigenomics over traditional bulk tissue assays.

More generally, our differential accessibility analysis was able to identify new enhancers by comparison with chromatin marks known to be associated with enhancers (Gjoneska et al. 2015). For example, when examining the most significantly differentially accessible loci for dentate granule cells, one of the top hits was a region marked by both H3K4me1 and H3K27ac, suggesting a putative enhancer upstream of the gene *Slc4a4* (Supplementary Fig. 7). *Slc4a4* encodes a sodium/bicarbonate co-transporter involved in mediating both intracellular and extracellular pH (Svichar et al. 2011), and *Slc4a4* expression is elevated in dentate granule neurons. While these accessible loci were enriched only in dentate neurons, several other accessible regions were identified in dentate granule cells as well as in the two pyramidal neuron populations, suggesting this gene is expressed in multiple cell types and, like *c-Fos*, may exhibit variable responses in different cell types.

### Cis regulatory networks in the hippocampus

Many enhancer elements are distal from the transcriptional start site of the gene. This results in a severe limitation in our ability to interpret the effects of epigenetic changes at these elements as they relate to gene expression. To address this limitation, groups have developed experimental proximity-ligation strategies, such as High throughput chromatin conformation capture (HiC; Lieberman-aiden et al. 2009), as well as computational strategies that associate regulatory elements with one another based on co-accessibility patterns. The latter approaches have typically relied on large databases of DNaseI hypersensitivity or ATAC-seq data performed on bulk cell isolates (Thurman et al. 2012; Budden et al. 2015); however, the same concepts have recently been applied to single-cell ATAC-seq data using the *Cicero* algorithm (Pliner et al. 2018). Briefly, *Cicero* uses an unsupervised machine-learning framework to link distal regulatory elements to their prospective genes via patterns of co-accessibility in the single-cell regulatory landscape. It achieves this by aggregating cells in close proximity and calculating a regularized correlation matrix between loci to produce co-accessibility scores between pairs or regulatory elements.

We applied *Cicero* to our hippocampus sci-ATAC-seq dataset to produce 487,156 links between ATAC-seq peaks at a co-accessibility score cutoff of 0.1 (Supplementary Data – InVivo.cicero_links.txt). Of these, 47,498 (10.5%) were links between two promoters, 146,818 (32.4%) linked a distal regulatory element to a promoter, and 259,236 (57.2%) were between two distal elements. We next compared our *Cicero*-linked peaks with existing chromatin conformation data that had been produced on mouse cortical tissue (Dixon et al. 2012), as no hippocampus data sets are currently available; however, a majority of topological associated domains (TADs) are conserved across cell types (Dixon et al. 2012). Consistent with expectations, we observed a 1.1 to 1.5 fold enrichment (Fig. 3A, *p* < 1×10^−4^ across all *Cicero* link thresholds out to 500 kbp, Methods) for linked peaks that occur within the same TAD over equidistant peaks present in different TADs, suggesting that the identified links are associated with higher-order chromatin structure. We then identified cis-co-accessibility networks (CCANs) using Cicero which employs a Louvain-based clustering algorithm, which can inform us about co-regulated chromatin hubs in the genome. Using a co-accessibility score threshold of 0.15 (based on high intra-TAD enrichment, Fig. 3A), we identified 3,243 CCANs, which incorporated 102,736 sites (mean 31.7 peaks/CCAN).

**Figure 3.**
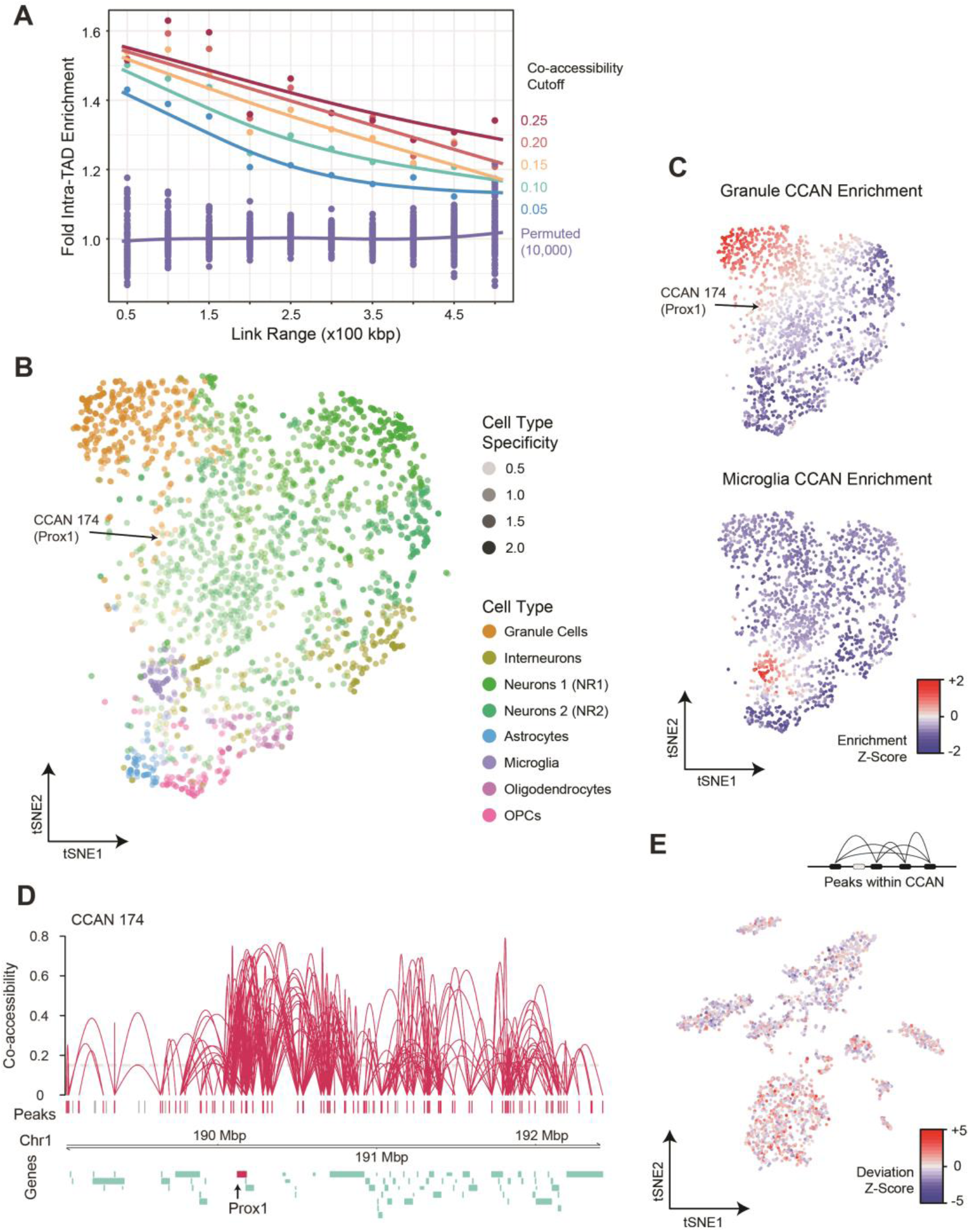
Cis co-accessibility analysis using *Cicero*. (A) *Cicero* links at several co-accessibility score thresholds are heavily enriched for links that contain peaks present in the same topological associated domain (TAD) as determined by Hi-C methods (Dixon et al. 2012). The enrichment decreases at greater distances (x-axis). (B) tSNE projection of CCANs colored by the cell type with the greatest accessibility for the CCAN. Each point represents an individual CCAN. Networks generally group by cell type. CCAN 174 which includes the *Prox1* gene shown below in (D) is indicated with an arrow. (C) Accessibility z-scores for CCANs for granule cells and microglia. (D) Cis co-accessibility network (CCAN) ID 174 including the *Prox1* promoter (dentate granule marker gene). (E) CCAN 174 has the greatest accessibility signal in cells identified as dentate granule cells.

To identify the enrichment of cell-type-specific CCANs, we first aggregated reads from cells within each cell type and calculated the fraction of cells that have signal within each peak of a CCAN. This assumes a uniform distribution of reads per cell across each cell type, which then allowed us to transform peak accessibility fractions across aggregate cell types into z-scores and then to average the signal within individual CCANs. The hierarchical clustering of the aggregate cell type populations reconstituted the relative clustering of the cell types (Supplementary Fig. 8). We further assessed cell type specific CCANs by projecting their relative similarity in two-dimensional tSNE space and visualized them based on their enrichment to their highest matching cell type (Fig. 3B,C, Supplementary Fig. 9). This analysis revealed distinct sets of co-accessibility networks for each cell type, with common networks falling towards the center of the projection space. CCANs with greater numbers of peaks tended to be less cell type specific, likely due to the large number of genes that are encompassed by the CCAN, the majority of which are not cell type specific (Supplementary Fig. 10). This observation is also consistent with chromatin conformation literature where common sets of topological domains are present across cell types with fewer that are cell type specific (Dixon et al. 2012) (Supplementary Fig. 11). Furthermore, CCANs also largely aggregated based on whether they are enriched in neuronal or non-neuronal cell types.

Included within our cell type specific CCANs, we observed a number of cell type marker genes. This included *Prox1*, a marker for dentate granule cells, which included 89 total accessibility sites and was associated with the correct population (Fig. 3D,E). While much of the CCAN did not exhibit cell type specificity, the region centered on Prox1 (with the highest co-accessibility values) drove the assignment. To dissect out the major components of the larger CCAN, we used *Cicero* specifically on the dentate granule cells (Supplementary Fig. 12A). This revealed three distinct CCANs within the *Prox1* region, with the central CCAN including the *Prox1* promoter, which exhibited the greatest specificity to the dentate granule cell cluster (Supplementary Fig. 12B). This suggests the possibility of larger chromatin networks with subsets of regulatory elements and genes joining or leaving the network based on cell type. Finally, we identified a number of CCANs that were overlapping that included mutually exclusive sets of peaks, suggesting two alternative folding patterns of chromatin within the regions dependent upon the cell type (Supplementary Fig. 13).

### In vitro neurons exhibit an altered epigenetic profile

To examine how well *in vitro* cultured hippocampal neuronal populations match their *in vivo* counterparts at the epigenetic level, we isolated hippocampal neurons from P0 pups and allowed them to mature for 16-18 days *in vitro* (DIV). At this stage, the neurons had extended long processes and expressed markers of mature neurons such as MAP2. We performed sci-ATAC-seq as described above and produced 899 high-quality single-cell chromatin accessibility profiles passing our quality thresholds (Methods). Our mean unique read count per cell was again high when compared to currently published work at 43,532, which to the best of our knowledge, is the highest achieved for any high-throughput single-cell ATAC-seq strategy to date. We then performed peak calling on the ensemble of *in vitro* sci-ATAC-seq profiles, resulting in 111,005 total peaks. Similar to our *in vivo* preparations, the ATAC-seq signal correlated well between the two replicates (Pearson R > 0.99). Subsequent filtering, LSI-tSNE, and clustering, as described for the *in vivo* preparation, revealed four distinct populations (Fig. 4A). Upon examination via marker gene and DNA binding motif accessibility enrichment, we determined one of the clusters to be the interneuron population (40.6% of cells), with the remainder being excitatory (59.4%).

**Figure 4.**
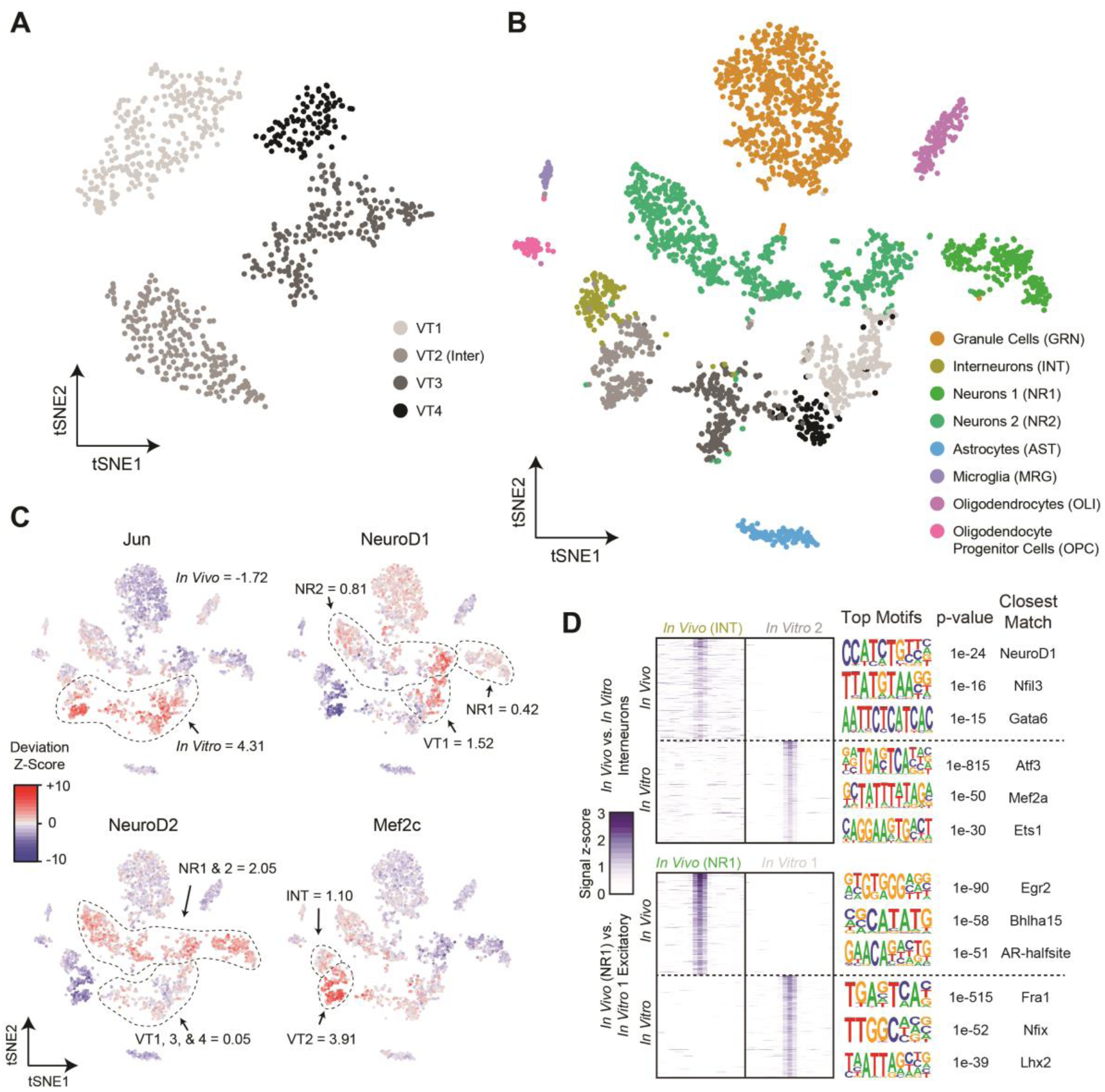
Comparison of the accessible chromatin landscape of *in vitro* cultured neurons with *in vivo* obtained profiles. (A) LSI-tSNE projections of *in vitro* obtained cells reveals four clusters, one of which exhibits interneuron patterns (VT2) and the remaining excitatory neurons (VT1,3-4). (B) LSI-tSNE projection of the combined *in vivo* and *in vitro* datasets colored by independently called clusters. Excitatory neurons in the two conditions generally cluster separately, with interneurons more closely associated. (C) *ChromVAR* global motif deviation z-scores for select motifs for each cell. Dashed lines and values correspond to mean values of cell populations. (D) Differential accessibility analysis between *in vivo* and *in vitro* interneurons (top, INT vs. VT2, respectively) and between two closest excitatory neuron populations between *in vivo* and *in vitro* conditions (NR1 and VT1, respectively). ATAC-seq signal is shown for the top differentially accessible loci with the top three motifs and corresponding *p*-values and matching motifs to the right.

We performed peak calling on the combined reads from both the *in vivo* and *in vitro* experiments and merged these peaks with those called on each set individually to produce a combined peak call set comprised of 174,503 sites. It is important to note that much of the increase over the *in vivo* peak set was due to increased coverage at sites that may not have met the calling threshold as opposed to peaks exclusive to the *in vitro* cultured neurons. We then performed LSI and tSNE on the resulting counts matrix using cells produced in both experiments. While the *in vitro* cultured glutamatergic neurons largely formed their own grouping independent of their *in vivo* counterparts, the inhibitory neurons from the *in vitro* preparation grouped more closely with the *in vivo* population (Fig. 4B).

We next examined the global DNA binding motif accessibility of the combined population (Fig. 4C). The starkest differences between the *in vivo* and *in vitro* cell populations was in motifs associated with the AP-1 complex, *i.e.* Fos, Jun, ATF, and JDP families (µz = 4.32 and −1.72 for *in vitro* and *in vivo* respectively). The AP-1 complex plays a major role in stimulus response, including cell stress (Hess 2004), which may not be surprising for neurons grown and matured *ex vivo*. It has also been shown that AP-1 modulates chromatin during neuronal activation (Su et al. 2017), suggesting the possibility of an elevated activity state in neuronal cultures compared to their *in vivo* counterparts; however, the decoupling of the many functional roles of the AP-1 complex from one another using global accessibility is not currently possible. We also examined the motifs for several other transcription factors that are relevant to neuronal development. NeuroD1, discussed above, responsible for early differentiation (E14.5 ventricular proliferative zone) (Pataskar et al. 2016) and survival of neurons, exhibited shared accessibility enrichment in a subset of cells from both the *in vivo* and *in vitro* neuronal populations. Mef2c delineates early precursors of a subset of inhibitory interneurons (Mayer et al. 2018) and we observed shared, elevated Mef2c accessibility in the interneuron populations, with greater accessibility in the *in vitro* cells (µ_z_ = 3.91) over that of the *in vivo* interneurons (µ_z_ = 1.10). In contrast to NeuroD1 and Mef2c, NeuroD2 acts later in hippocampal development than NeuroD1 (Pleasure et al. 2000), is expressed in migrating granule neurons, and binds to a number of neuron-specific promoters. The DNA binding motif for NeuroD2 was globally more accessible in the *in vivo* neurons when compared to their *in vitro* counterpart (µ_z_ = 2.05 and µ_z_ = 0.05 for *in vivo and in vitro* respectively). This finding may reflect its later developmental appearance and that the main targets of NeuroD2 are involved in layer-specific differentiation and axonal pathfinding, which are not likely to be occurring *in vitro*.

Differential accessibility analysis comparing *in vitro* and *in vivo* counterparts shed further light on the epigenetic differences between the two populations (Fig. 4D). A comparison of the interneuron populations produced 4,356 and 7,575 peaks significantly differentially accessible in the *in vivo* (INT) and *in vitro* (VT2) populations, respectively (*q*-value ≤ 0.01, Log2 fold-change ≥ 1). A motif enrichment analysis of these peak sets revealed the most significantly enriched motifs corresponded to NeuroD1 in the *in vivo* peaks (*p* = 1×10^−24^), which is interesting because NeuroD1 global accessibility is low in both interneuron populations (Fig. 4C). Interneuron peaks specific to the *in vitro* population were significantly enriched for Atf3 (*p* = 1×10^−815^), which is not surprising in light of the above accessibility of AP-1 in the *in vitro* cell populations and its shared role in cell stress and interaction with the AP-1 complex (Hai and Curran 1991). We also examined differential accessibility between the most-closely grouped excitatory neuronal populations, which produced 1,761 and 2,964 for NR1 (*in vivo*) and VT1 (*in vitro*) respectively (*q*-value ≤ 0.01, Log_2_ fold-change ≥ 1). The most significantly enriched motif in the *in vivo* peak set was Egr2 (*p* = 1×10^−90^), again a transcription factor expressed highly in migrating neural crest cells (Wilkinson et al. 1989) that may be absent in an *in vitro* setting where cell migration is not pertinent.

## Discussion

A better understanding of the role of specific cell populations in hippocampal function is a necessary step in order to study disease processes that involve this region critical to memory and learning. Thus far, studies have used gene expression data from sorted populations (Cembrowski et al. 2016) and single cells (Zeisel et al. 2015; Habib et al. 2017) to identify subpopulations of cells and novel marker genes for the cells within the hippocampus. However, gene expression data is typically graded across cell types with few cases of exclusive expression, making it difficult to detect significant change within a cell type. In contrast, chromatin accessibility is typically a binary signature and involves far more regulatory elements than there are genes, thus motivating the use of ATAC-seq methods for cell type deconvolution and assessment.

Here, we provide the most in-depth epigenetic analysis of the hippocampus at single-cell resolution to date. Our sci-ATAC-seq protocol (Methods) has been optimized for primary cell culture and both fresh or frozen tissue and produces unique read counts per cell in the tens-of-thousands, a full order-of-magnitude improvement over the initial sci-ATAC-seq publication (Cusanovich et al. 2015). The data sets released with this study can be readily analyzed using *scitools* (https://github.com/adeylab) to recreate the figures in this manuscript in addition to many more, such as the gamut of motif accessibilities that we assessed. This tool suite is designed to be complementary to other single-cell ATAC-seq analysis packages, such as *ChromVAR* and *Cicero*, and serves as an easy framework for integrating analyses and generating plots to assess data quality and facilitate biological interpretation. *Scitools* is also set up to directly feed sci-ATAC-seq data into *Monocle3* for pseudotemporal ordering applications not described in this work.

We utilized our sci-ATAC-seq maps to identify the major cell types of the hippocampus, with sufficient depth and library complexity to profile less abundant cell types, such as microglia and oligodendrocyte progenitor cells. Our analysis of global motif accessibility revealed the expected enrichment of motifs associated with specific cell populations in addition to uncovering unanticipated findings, such as increased accessibility at CTCF motifs in interneuron and glial populations; a finding that was also observed in our differentially accessibility analysis. We also utilized our dataset to map cis co-accessibility networks, enabling the association of distal elements with promoters or other regulatory loci. Finally, we directly compared the accessibility profiles of neurons that were matured *in vitro* with their *in vivo* counterparts. This revealed a stark difference in the global accessibility for motifs associated with the AP-1 complex, which is involved in cell stress as well as neuronal activity. Future work to identify the cause and effect of elevated AP-1 complex activity is warranted to understand its impact on studies that utilize hippocampal neurons matured *in vitro*.

We believe that the chromatin accessibility maps we provide in this work, including the profiling of *in vitro* cultured neurons, and the software tools we are releasing are a valuable resource for any groups studying the hippocampus or those that wish to analyze single-cell chromatin accessibility data.

## Methods

### Isolation of hippocampus tissue

All animal studies were approved by the Oregon Health and Science University Institutional Animal Care and Use Committee. Sixty day old C57BL/6J mice were deeply anesthetized using isofluorane. After decapitation the brain was removed and the hippocampus isolated and placed in ice-cold phosphate-buffered saline (pH 7.4).

### In Vitro culturing of hippocampal neurons

Pups (P0) were killed by decapitation and the brains dissected in ice-cold Hanks Basal Salt Solution (HBSS, pH 7.4) with 25 mM Hepes buffer. Individual hippocampi were excised without the meninges and pooled by individual animal. The tissue was treated with 2% papain and 80ng/ml Dnase I in HBSS at 37 °C for 10 min. Tissue pieces were rinsed three times with Hibernate A containing 2mM Glutamax and 1× B27 supplement. Neurons were dissociated carefully and filtered with a 0.4-μm mesh. Neurons were plated at a density of 1×10^6^ cells per well of a six well dish coated with 50 μg/mL Poly-L-Lysine hydrobromide in boric acid buffer (50 mM Boric Acid, 12.5 mM Sodium Borate, decahydrate). The neurons were plated in Neurobasal A containing 1xB27 supplement and 2mM glutamax. After 2 hours, the media was changed to remove cell debris. Media half changes occurred every 3 days with fresh Neurobasal A containing 1xB27 and 2mM glutamax. Cells were maintained at 37°C with 5% CO_2_ in a humidified incubator.

### Sci-ATAC-seq assay & sequencing

Tissue was diced on ice using a sterile razor blade in freshly-prepared Nuclei Isolation Buffer (NIB: 500 µL 10 mM Tris-HCl pH.5, 100 µL 10 mM NaCl, 150 µL MgCl2, 500 µL 0.1% Igepal, 1 unit Qiagen Protease Inhibitor, nuclease-free water to 50 mL) followed by dounce homogenization. For cultured cells, nculei were directly isolated by removing media, washing once with ice cold PBS, and then NIB added to cover the dish followed by incubation on ice for 5 minutes, scraping using a tissue scraper, and then an additional 5-minute incubation on ice. For both tissue and cultured cells, nuclei were then pelleted and resuspended in 1 mL NIB with DAPI added to a final concentration of 5 mg/mL. Nuclei were then strained in a 35 µm strainer and sorted on a Sony SH800 Flow Sorter and deposited into 0.2 mL PCR plates containing 5 uL of 2X TD buffer and 5 uL of NIB, with 2,500 nuclei deposited per well. Plates were placed on ice until transposition. Tagmentation was performed by the addition of 1 µL of 2.5 µM barcoded transposome (Amini et al. 2014) and incubated at 55°C for 15 minutes followed by placing the plate on ice to stop the reaction. All wells were then pooled using wide-bore pipette tips and DAPI added to a final concentration of 5 mg/mL. Tagmented nuclei were then strained and sorted again and 22 were deposited into each new PCR well containing 0.25 µL 20 mg/mL BSA, 0.5 µL 1% SDS, 7.75 µL nuclease-free water, 2.5 µL barcoded forward primer, and 2.5 µL reverse primer. Plates were kept on ice until all sorting was completed. After sorting, plates were incubated at 55°C for 15 minutes to denature the transposase followed by placing the plate on ice and adding 12 uL of PCR mix (7.5 µL NPM, 4 µL nuclease-free water, 0.5 µL 100X SYBR Green) and then PCR amplified using the following conditions: 72°C for 5:00, 98°C for 0:30, Cycles of [98°C for 0:10, 63°C for 0:30, 72°C for 1:00, plate read, 72°C for 0:10] on a BioRad CFX real time thermocycler. Reactions were pulled when mid-exponential, typically 17-22 cycles. Post-amplification 5 µL of each reaction was pooled and cleaned up using a Qiaquick PCR Purification column. Libraries were quantified using a Qubit fluorimeter, diluted to ∼4 ng/µL and assessed on an Agilent Bioanalyzer HS Chip. Sequencing was carried out as previously described on a NextSeq^™^ 500 (research use only) using custom primers and chemistry (Vitak et al. 2017).

### The scitools suite

All initial analysis was performed with *scitools,* a custom software package we developed to help analyze sci-ATAC-seq data and other combinatorial indexing data (sci-). The toolset is a collection of commands to perform common functions for sci-datasets, including wrappers that utilize existing tools, including: *bwa* (Li and Durbin 2009), *macs2* (Zhang et al. 2008), *bedtools* (Quinlan and Hall 2010), *samtools*, as well as R libraries: *ggplot2, chromVAR* (Schep et al. 2017), *chromVARmotifs, cicero* (Pliner et al. 2018), *RtSNE, dbscan* (Ester et al. 1996). Usage of *scitools* for any of these functions should cite the relevant utilities. Data produced for this study are available on GEO under accession GSE118987 as reads as well as data tables and metadata in the form of data.gz files which can be split into their components via “scitools split-data”. See Supplementary Note 1 document and Supplementary Figure 14 for additional details on datasets provided as well as the scitools documentation, which can be found at https://github.com/adeylab/scitools.

### Sci-ATAC-seq data processing

BCL files were first converted to fastq files using bcl2fastq (2.19.0). We then demultiplexed our reads using scitools (fastq-dump, fastq-split) based on the two separate Tn5 tagmentation events on the P5 and P7 ends of the molecules and the following added unique PCR indexes on both sides. In order for a barcode to be considered a match each of these four indexes constituting a barcode had to be within two Hamming edit distances away from their expected counterpart. We aligned to the mm10 genome using the scitools fastq-align function within scitools, which mapped reads using *bwa mem*. Aligned reads were filtered based on a quality score cutoff of 10 and PCR duplicates removed in a barcode-aware manner using scitools bam-rmdup. We determined whether a barcode represented a cell as opposed it representing noise by using the mixed model approach previously presented (Vitak et al. 2017). Peaks were then called using scitools callpeak, which utilizes macs2 to identify peaks and then extend to 500 bp followed by peak merging and filtering of peaks that extend beyond chromosome boundaries.

### Latent semantic indexing and 2D embedding

Count matrixes were generated using scitools counts to produce a matrix of read counts at cells (columns) by called peaks (rows). This matrix was then filtered using scitools filter-matrix to exclude rows with fewer than 10 cells having reads (-R 10), and columns (cells) with fewer than 1000 rows with reads (-C 1000). The matrix was then carried through term-frequency inverse-document-frequency transformation using scitools tfidf, followed by latent semantic indexing, retaining SVD dimensions 1-15 using scitools lsi. The resulting LSI matrix was used in scitools tsne which makes use of the *RtSNE* R package. All tSNE plots were generated using scitools plot-dims using an annotation file to encode cluster ID, sample ID, or other variables, including *chromVAR* motif deviation z-scores.

### Identifying transcription-facter-associated changes

We applied the *chromVAR* (Schep et al. 2017) R package to our data to infer changes in global motif accessibility across our cell populations. This provides information on the putative binding of transcription-factors and consequently the possible ongoing biological processes in cell populations. The mouse_pwms_v1 motif set from the *chromVARmotifs R* package was used in this analysis. The bias corrected motif deviation scores were plotted on the tSNE embedded 2D coordinates with the *scitools* plot-dims -M option for visualization.

### Cell type dependent differential accessibility

To accurately identify differentially accessible peaks we used the *make_glasso_cds* function from the *Cicero* (ver=,0.0.0.9000) package to create clusters of k=50 cells based on their the low dimensional t-SNE coordinates. We then selected clusters with 99% cell type purity and aggregated accessibility profiles. We posited that the aggregate profiles would provide the replicates required for the DEseq2 R package, which in turn internally corrects for technical biases such as assay efficiency. With this method we tested (using the inherent *nBinomWaldTest*) for differentially accessible sites between cell types against all other cell types combined. We corrected for multiple testing at q=0.01 and further filtered differentially accessible sites by removing peaks accessible at q=0.2 in any of the other cell types. We also note that scitools aggregate-cells is also capable of aggregating cells in reduced dimensional space for purposes of differential accessibility analysis. We then applied Homer (http://homer.ucsd.edu/homer/motif/) to identify potential *de novo* and known regulators of chromatin accessibility within the cell type dependent differentially accessible sites. We used all accessible peaks as background and the mm10 findMotifsGenome command.

### Identifying cis-regulatory networks in the hippocampus

We used the recently described *Cicero* package (Pliner et al. 2018) to identify cis-co-accessibility networks (CCANs). This method uses the patterns of co-accessibility in sci-ATAC-seq data to link distant regulatory elements with their target genes. It achieves this by first grouping cells together into local neighborhoods based on their coordinates in low dimensional t-SNE space and creating aggregate accessibility profiles from their single-cell chromatin accessibility matrixes. The aggregate counts are then bias corrected for technical factors and a raw site-to-site covariance matrix is calculated for overlapping windows across the genome. At this point the algorithm uses a graphical LASSO model, to calculate a regularized correlation matrix that applies increasing penalty based on the genomic distance between two sites. Finally, the overlapping covariance matrixes are collapsed resulting in a matrix of co-accessibility scores that links sites across cell neighborhoods. CCANs are constructed using the Louvain community detection algorithm that creates subgraphs from the network of linked sites. This is done based on the values from the co-accessibility score matrix that are above a given threshold. We used a p=0.15 threshold cutoff for and identified 2,066 chromatin networks which incorporated 47,805 sites of our *in vivo* cell populations.

Fold enrichment for links within annotated TADs (Dixon et al. 2012) was performed by calculating the proportion of distance-matched (±25 kbp of specified 50 kbp distance interval) intra-TAD links over inter-TAD links at a range of co-accessibility score cutoffs (0.05 to 0.25 at 0.05 intervals). 10,000 permutations were then performed for each distance bin by randomly assigning two distance-matched peaks as linked and retaining the same total number of links for each co-accessibility cutoff and then calculating the fold intra-TAD enrichment as described above.

### Cell type specific cis-regulatory networks

We aimed to assign cis-co-accessibility networks to cell types using their relative accessibility across these groups. We approached this by first calculating the fraction of cells of each cell type that have signal at a peak. We then assumed that the distribution of reads per cell across these cell types is close to uniform, which allowed us to aggregate the z-scored read fractions of peaks within our CCANs. We finally z-scored the resulting matrix across the CCANs and then visualized the separation of CCANs by cell type by bi-clustering and plotting the heatmap using the *complexHeatmap* (ver=1.17.1) R package. We also visualized CCAN cell type specificity by using tSNE on the z-scored group read fractions to embed CCANs in 2D. We assigned the cell type to each of the CCANs based on the highest z-scored value. We next identified CCANs that contain at least one of the genes (*Prox1, Dsp, Ociad2, Dkk3, Glul, Gfap, Mog, Cldn11, C1qa, Wfs1, Mobp, Pdgfra*) shown to be differentially accessible in our data. We intersected +/-80 kbp regions before and after transcription start sites of these genes with the CCANs using bedtools intersect. We plotted the CCANs around genes where the cell type assigned to the CCANs matched the cell type specificity of the gene using the cicero *plot_connections* function. We used *chromVAR* to further validate the relative enrichment of CCANs by using CCAN peaks as motif input files. We used *scitools* plot dims -M option to visualize the deviation scores for the CCANs on the tSNE coordinates. We have to note that in order for this method to work, peaks within the CCANs had to be accessible across multiple cell types, so we decided to use only CCANs with ≥ 10 peaks for this analysis. We finally included a more in-depth analysis of CCAN 174 centered around *Prox1*. We called CCANs just within Granule cells and identified three different sub CCANs, with the core of the original CCAN 174 showing even higher specify in the *chromVAR* deviation scores plots (Supplementary Fig. 11).

## Acknowledgements

We would like to thank Ryan Mulqueen and Andy Fields for helpful suggestions and contributions along with other members of the Adey Lab. This work was supported by NIH (R35GM124704 to A.C.A., NS099374 to G.M., U54DK107979 and DP2HD088158 to C.T.), an NSF GRFP to H.A.P.

(DGE-1256082), and support from the Rett Syndrome Research Trust to G.M.

## Author Contributions

A.C.A., J.R.S., and G.M. designed all experiments. A.C.A., J.R.S., K.A.T., and M.W.L. wrote the manuscript. S.A.V. performed all sci-ATAC-seq preparations. A.C.A. and K.A.T. performed computational analysis and wrote software associated with this work. J.R.S. prepared all tissue samples and cultures, and performed analyses and interpretation of the data. M.W.L. aided in analysis of the data and interpretation of findings. H.A.P. and C.T. provided early access to computational tools and aided in the co-accessibility analysis and interpretation. F.J.S. provided reagents and contributed to sci-ATAC-seq method development and implementation. All authors reviewed and approved the manuscript.

## Supplementary Information

**Supplementary Table 1:**
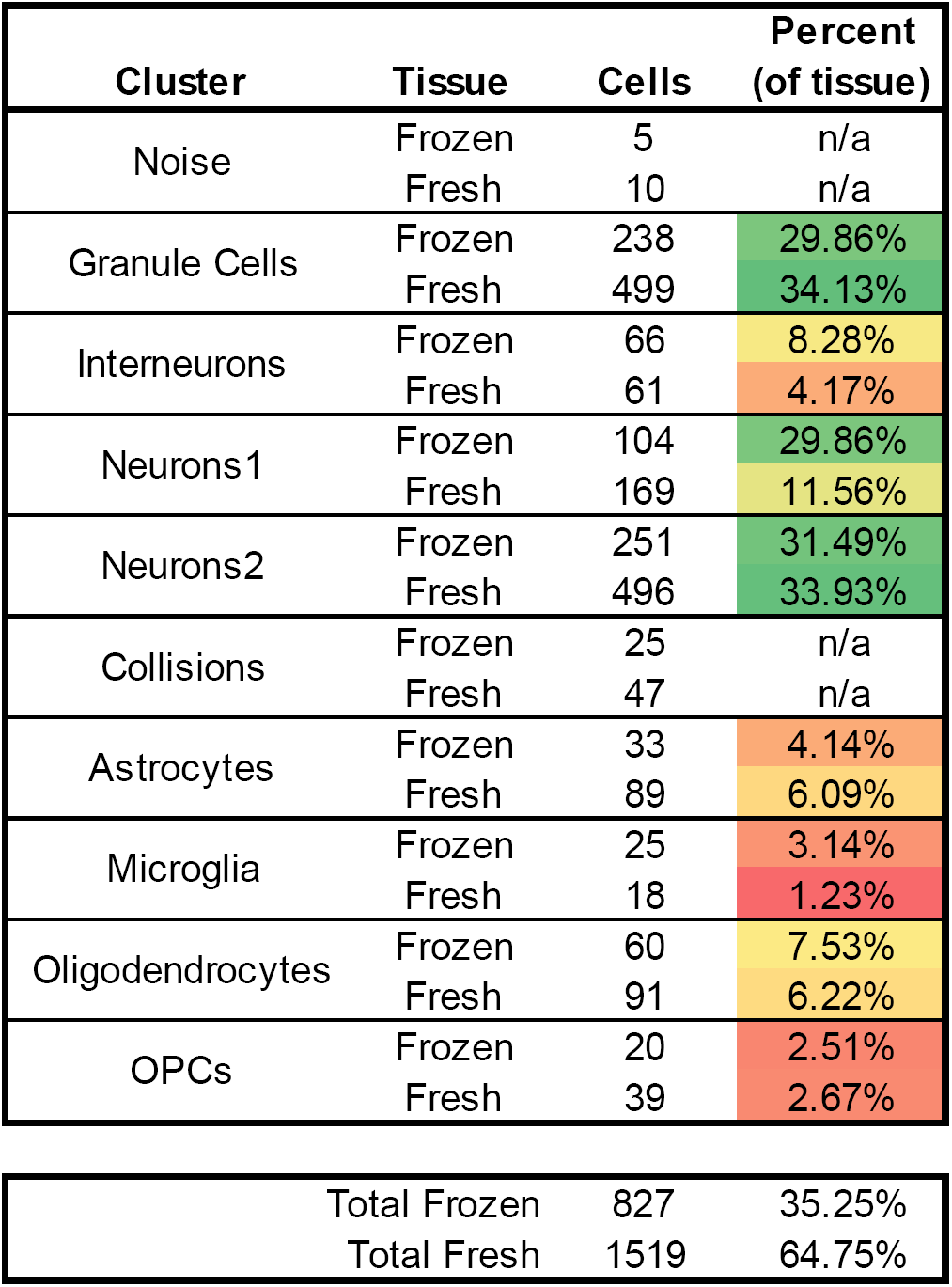
Cell Type Composition Proportion of cells assigned to each cluster for the *In Vivo* dataset along with the fresh vs. frozen breakdown.

| Cluster | Tissue | Cells | Percent<br>(of tissue) |
| --- | --- | --- | --- |
| Noise | Frozen | 5 | n/a |
|  | Fresh | 10 | n/a |
| Granule Cells | Frozen | 238 | 29.86% |
|  | Fresh | 499 | 34.13% |
| Interneurons | Frozen | 66 | 8.28% |
|  | Fresh | 61 | 4.17% |
| Neurons1 | Frozen | 104 | 29.86% |
|  | Fresh | 169 | 11.56% |
| Neurons2 | Frozen | 251 | 31.49% |
|  | Fresh | 496 | 33.93% |
| Collisions | Frozen | 25 | n/a |
|  | Fresh | 47 | n/a |
| Astrocytes | Frozen | 33 | 4.14% |
|  | Fresh | 89 | 6.09% |
| Microglia | Frozen | 25 | 3.14% |
|  | Fresh | 18 | 1.23% |
| Oligodendrocytes | Frozen | 60 | 7.53% |
|  | Fresh | 91 | 6.22% |
| OPCs | Frozen | 20 | 2.51% |
|  | Fresh | 39 | 2.67% |
| Total Frozen |  | 827 | 35.25% |
| Total Fresh |  | 1519 | 64.75% |

**Supplementary Table 2:** Differentially Accessible Peaks The top 10 differentially accessible (see Methods) peaks corresponding to each cluster. Plots for these can be found in Supplementary Fig. 4.

|  | Chrom. | Start | End |  | Chrom. | Start | End |  | Chrom. | Start | End |
| --- | --- | --- | --- | --- | --- | --- | --- | --- | --- | --- | --- |
| Granule Cells | chr2 | 12,805,030 | 12,805,581 | Neurons 2 (CA3) | chr19 | 23,330,204 | 23,331,209 | Oligodendrocytes | chr10 | 83,906,604 | 83,907,477 |
|  | chr18 | 13,157,877 | 13,158,511 |  | chr2 | 169,049,869 | 169,050,512 |  | chr12 | 100,073,051 | 100,073,764 |
|  | chr19 | 24,665,688 | 24,666,194 |  | chr15 | 94,011,340 | 94,012,637 |  | chr18 | 75,544,174 | 75,544,677 |
|  | chr8 | 24,303,072 | 24,303,600 |  | chr19 | 18,368,387 | 18,369,139 |  | chr13 | 56,581,977 | 56,583,211 |
|  | chr14 | 11,231,939 | 11,232,458 |  | chr9 | 97,883,149 | 97,883,765 |  | chr18 | 75,541,370 | 75,541,889 |
|  | chr1 | 190,306,260 | 190,306,838 |  | chr6 | 55,836,835 | 55,837,355 |  | chr9 | 25,230,456 | 25,231,009 |
|  | chr1 | 190,642,627 | 190,643,179 |  | chr18 | 78,243,805 | 78,244,339 |  | chr17 | 8,350,898 | 8,351,641 |
|  | chr5 | 88,830,139 | 88,830,687 |  | chr3 | 65,884,483 | 65,885,067 |  | chr2 | 127,627,327 | 127,628,521 |
|  | chr1 | 55,828,229 | 55,828,748 |  | chr10 | 111,308,849 | 111,309,852 |  | chr14 | 34,590,699 | 34,591,715 |
|  | chr10 | 48,010,760 | 48,012,230 |  | chr1 | 127,541,771 | 127,542,290 |  | chr9 | 25,240,906 | 25,241,521 |
| Interneurons | chr2 | 158,610,479 | 158,611,273 | Astrocytes | chr3 | 50,421,595 | 50,422,393 | Oligo. Progenitors | chr5 | 65,434,477 | 65,435,230 |
|  | chr2 | 71,529,369 | 71,529,901 |  | chr10 | 18,359,980 | 18,360,766 |  | chr2 | 125,723,709 | 125,725,049 |
|  | chr11 | 88,028,257 | 88,028,867 |  | chr2 | 102,620,556 | 102,621,235 |  | chr9 | 56,868,193 | 56,868,694 |
|  | chr2 | 22,621,845 | 22,622,763 |  | chr14 | 65,889,584 | 65,890,606 |  | chr6 | 30,572,647 | 30,573,201 |
|  | chr6 | 55,398,876 | 55,399,381 |  | chr14 | 121,113,651 | 121,114,302 |  | chr7 | 79,729,178 | 79,729,709 |
|  | chr5 | 124,206,256 | 124,206,918 |  | chr1 | 154,139,439 | 154,140,079 |  | chr14 | 57,534,993 | 57,535,499 |
|  | chr5 | 31,589,699 | 31,590,705 |  | chr16 | 18,083,199 | 18,083,957 |  | chr4 | 106,317,513 | 106,318,014 |
|  | chr2 | 36,011,161 | 36,011,981 |  | chr1 | 182,483,531 | 182,484,443 |  | chr3 | 51,749,422 | 51,750,382 |
|  | chr5 | 35,985,506 | 35,986,014 |  | chr5 | 33,705,059 | 33,705,607 |  | chr1 | 172,064,528 | 172,065,243 |
|  | chr19 | 15,052,684 | 15,053,503 |  | chr12 | 25,239,950 | 25,241,190 |  | chr6 | 146,955,683 | 146,956,269 |
| Neurons 1 (CA1) | chr18 | 15,008,128 | 15,008,636 | Microglia | chr7 | 67,445,504 | 67,446,168 |  |  |  |  |
|  | chr9 | 22,030,329 | 22,030,831 |  | chr7 | 126,746,073 | 126,747,014 |  |  |  |  |
|  | chr8 | 40,455,798 | 40,456,317 |  | chr15 | 27,702,515 | 27,703,332 |  |  |  |  |
|  | chr7 | 84,576,178 | 84,576,680 |  | chr7 | 80,078,688 | 80,079,367 |  |  |  |  |
|  | chr13 | 73,912,658 | 73,913,254 |  | chr15 | 59,728,185 | 59,728,766 |  |  |  |  |
|  | chr2 | 46,330,878 | 46,331,408 |  | chr7 | 145,111,444 | 145,112,140 |  |  |  |  |
|  | chrX | 145,341,895 | 145,342,781 |  | chr4 | 129,782,344 | 129,783,082 |  |  |  |  |
|  | chr6 | 89,050,828 | 89,051,367 |  | chr13 | 83,571,978 | 83,572,659 |  |  |  |  |
|  | chr10 | 54,551,634 | 54,552,521 |  | chr17 | 88,581,421 | 88,581,940 |  |  |  |  |
|  | chr3 | 74,734,781 | 74,735,299 |  | chr7 | 120,851,508 | 120,852,400 |  |  |  |  |

**Supplementary Figure 1:**
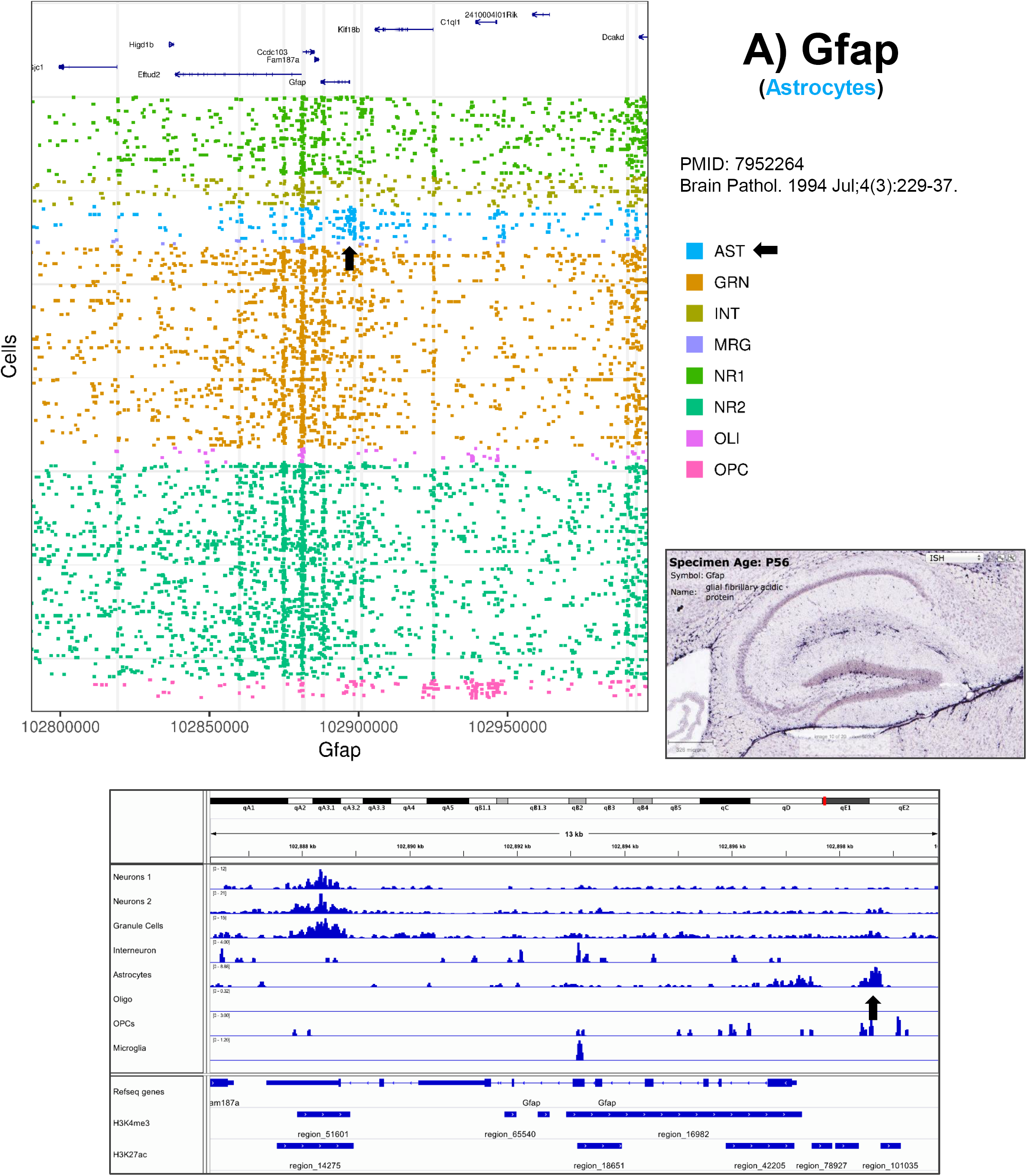
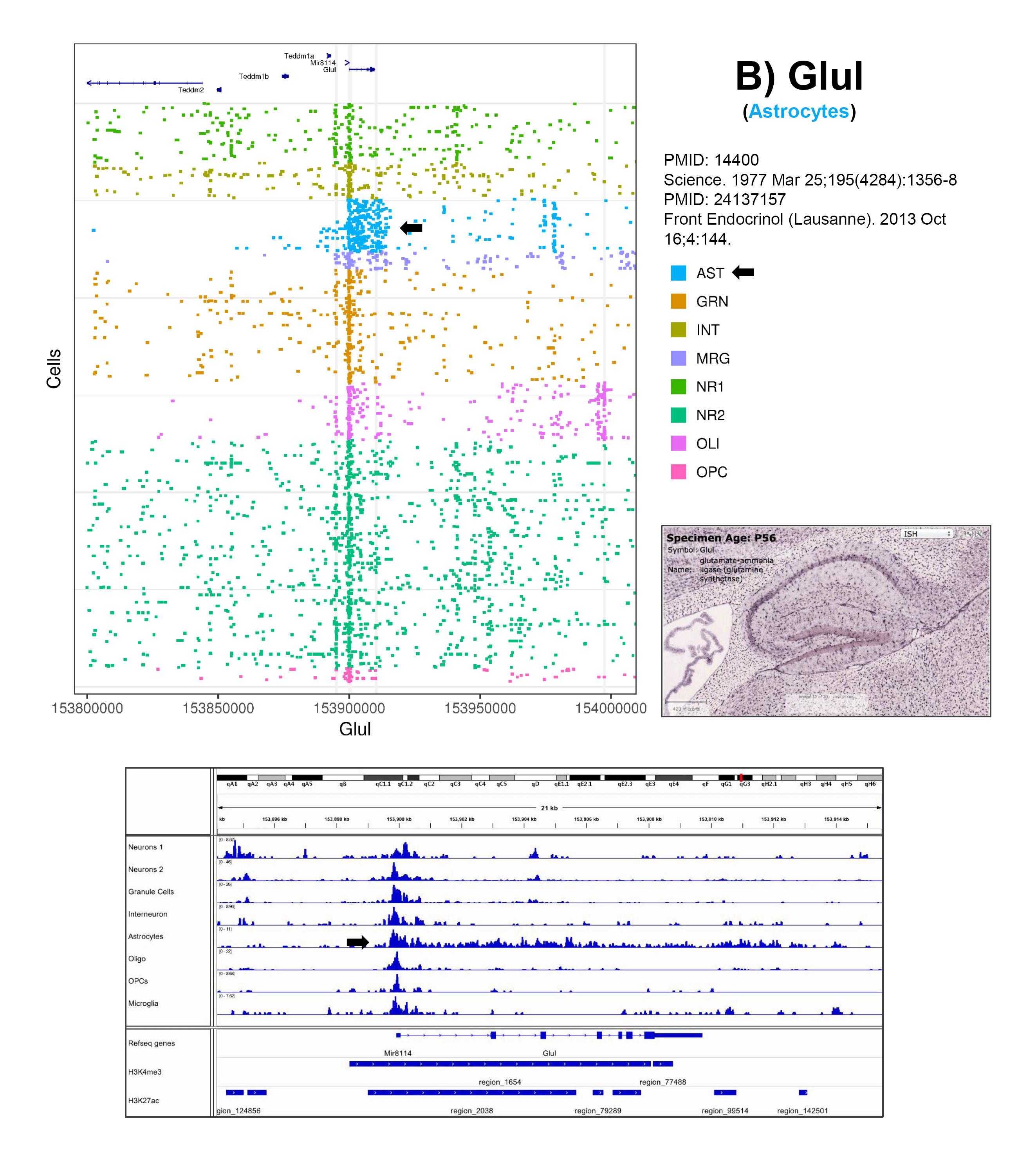
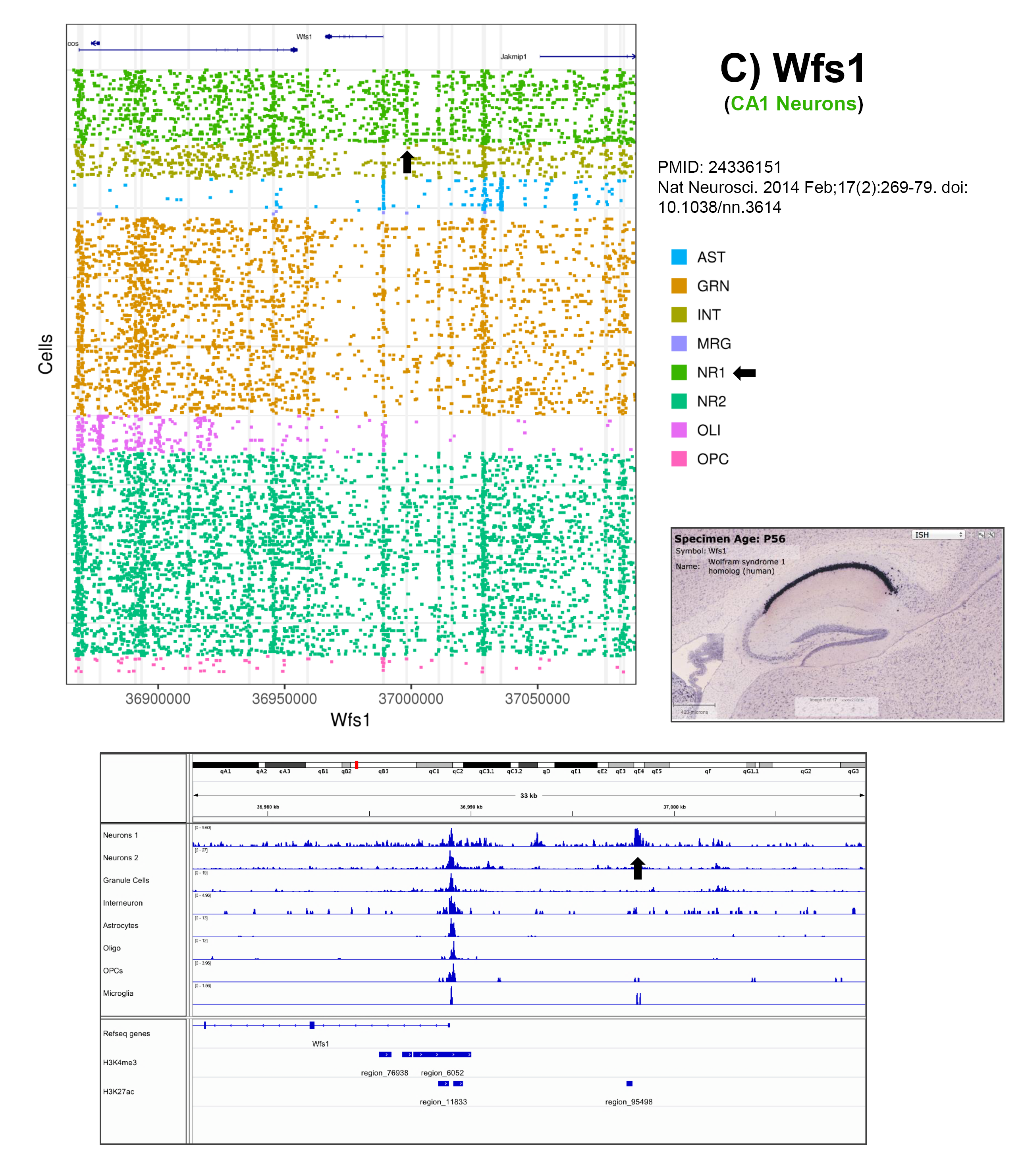
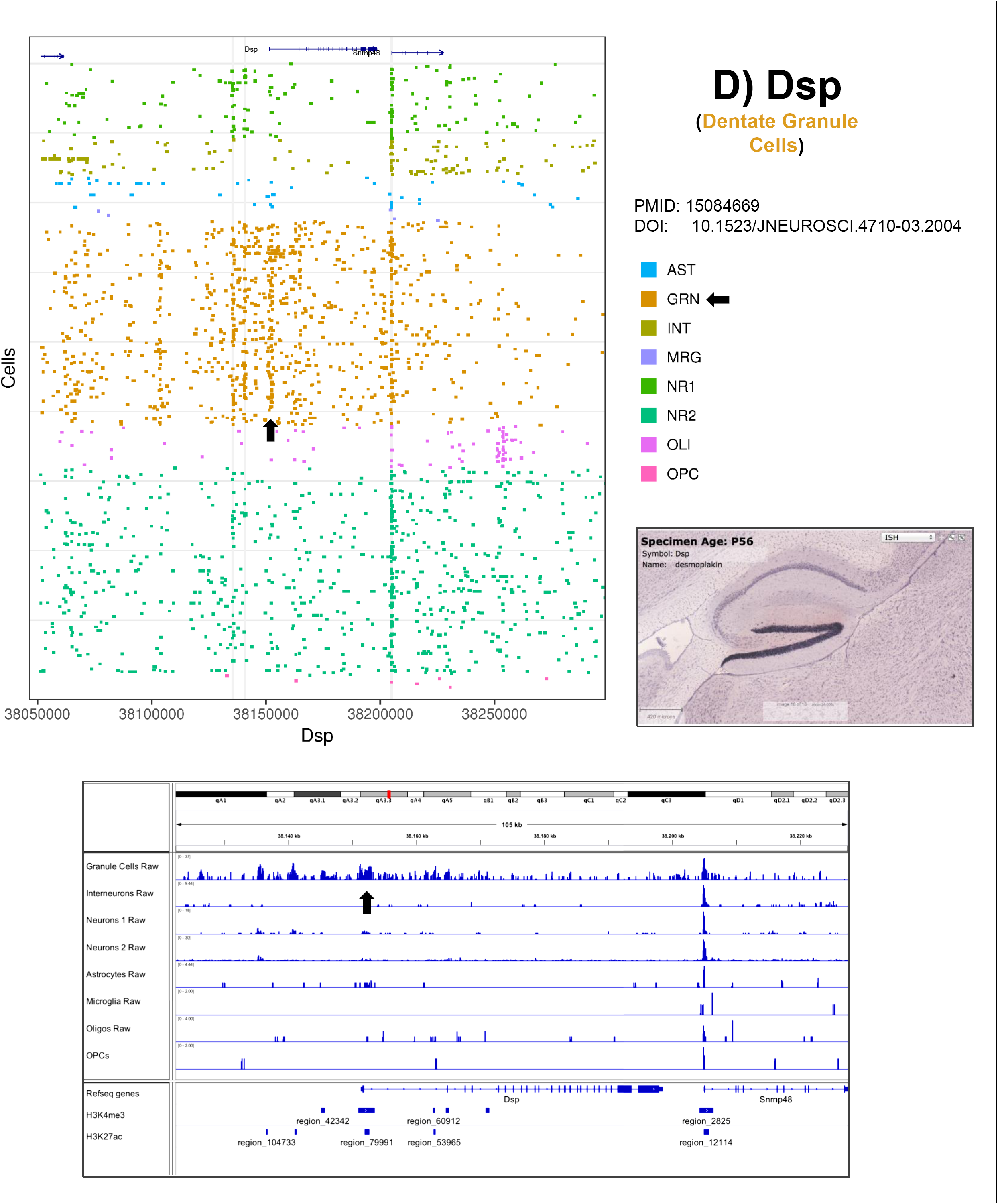
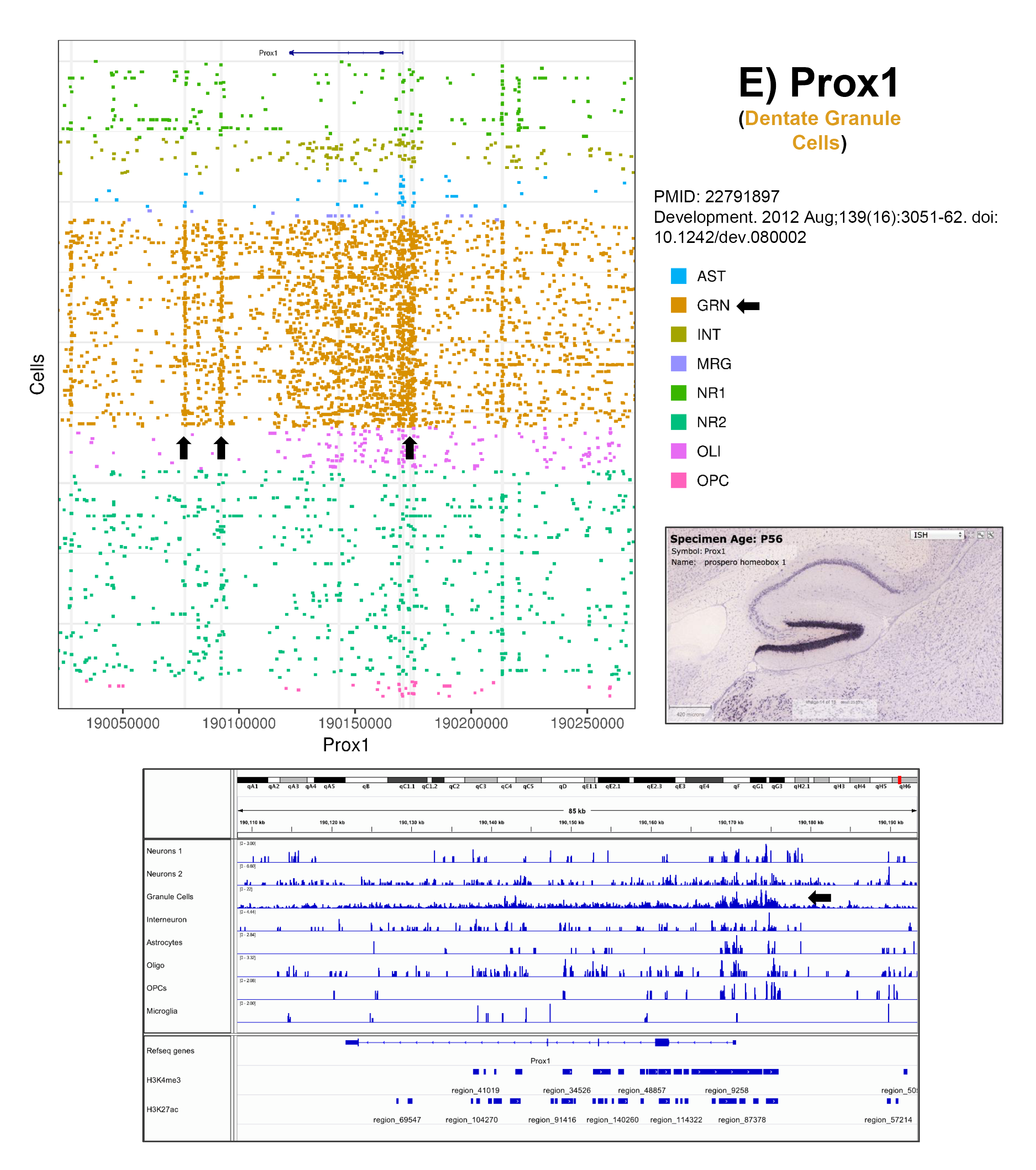
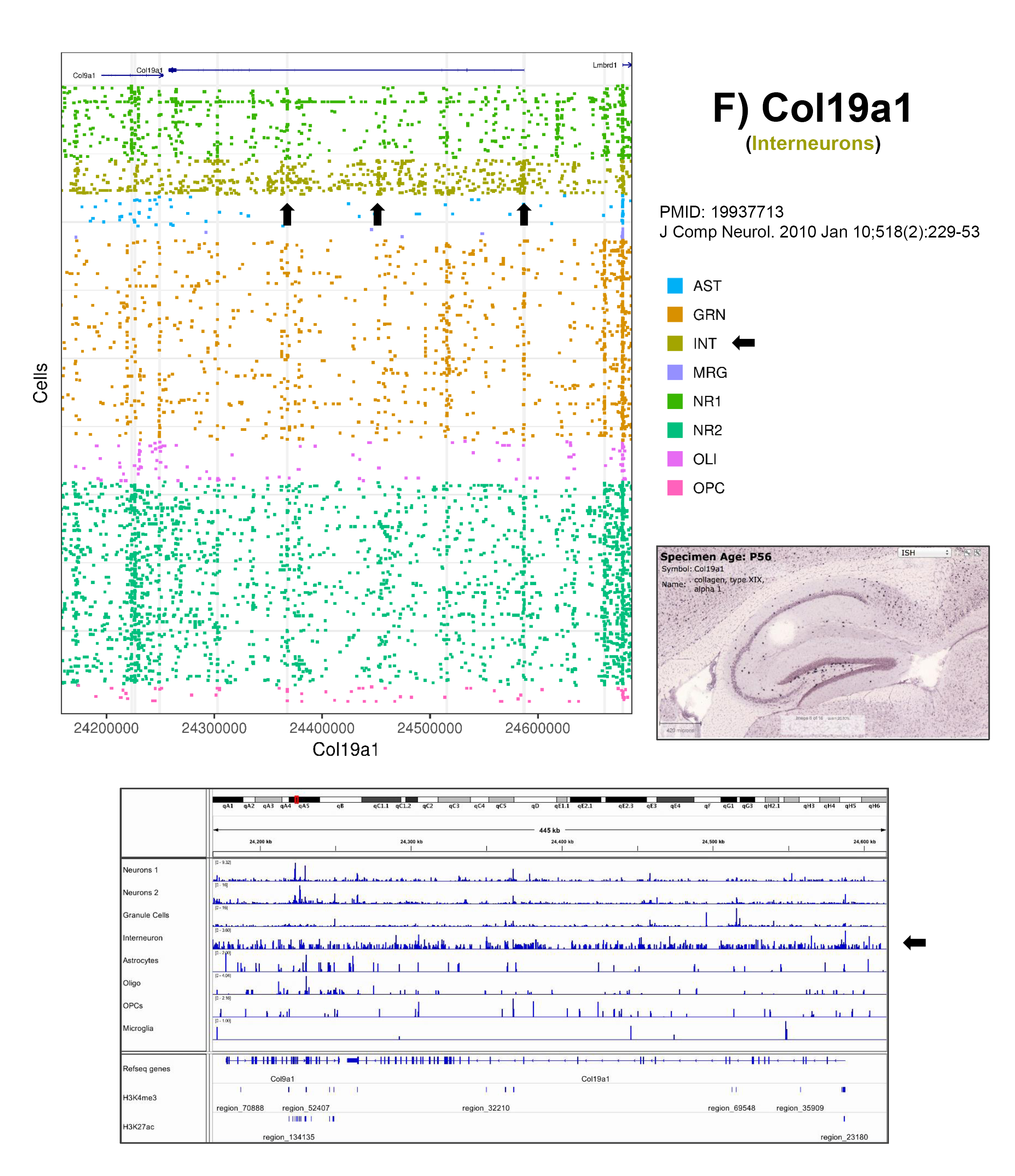
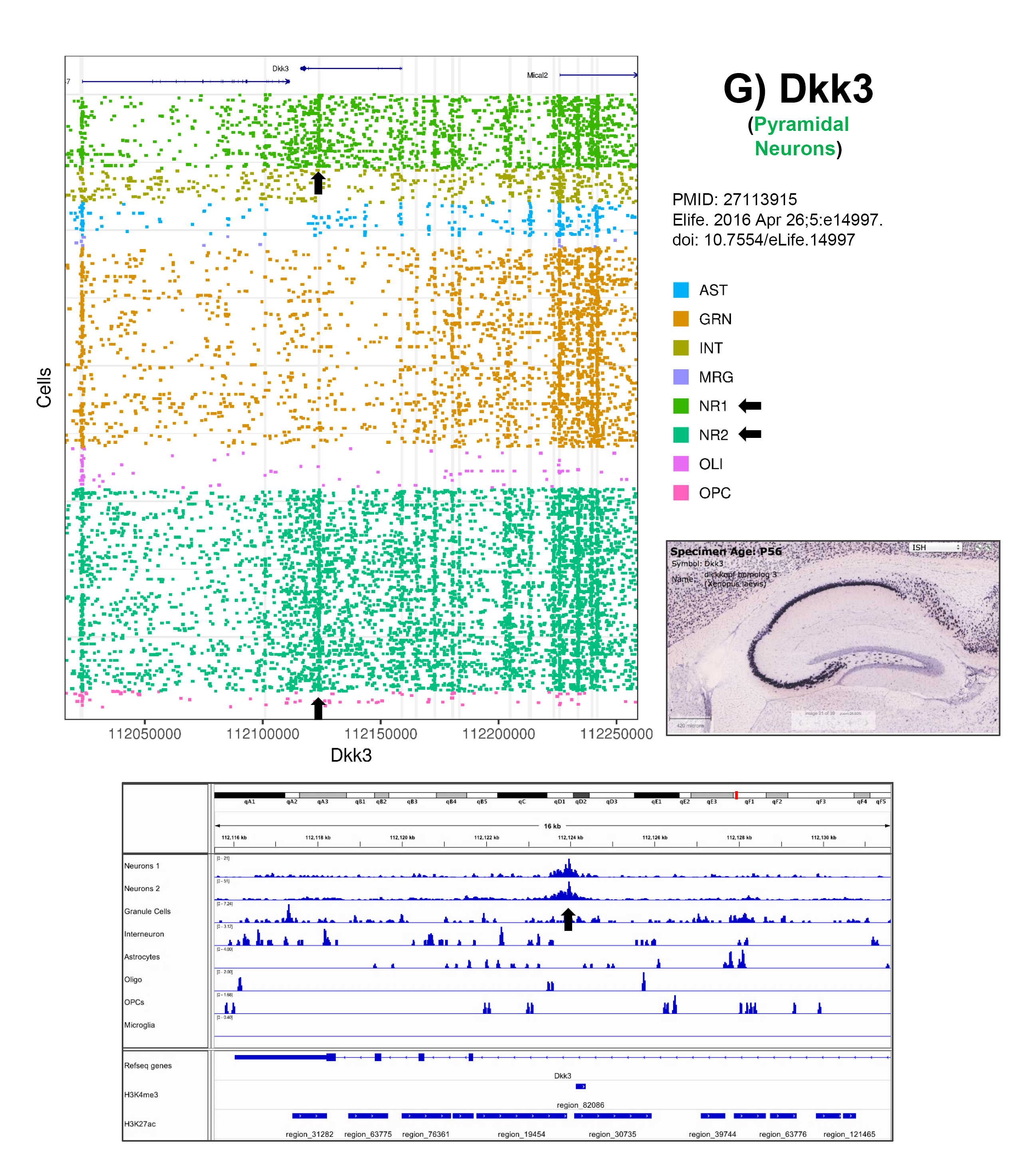
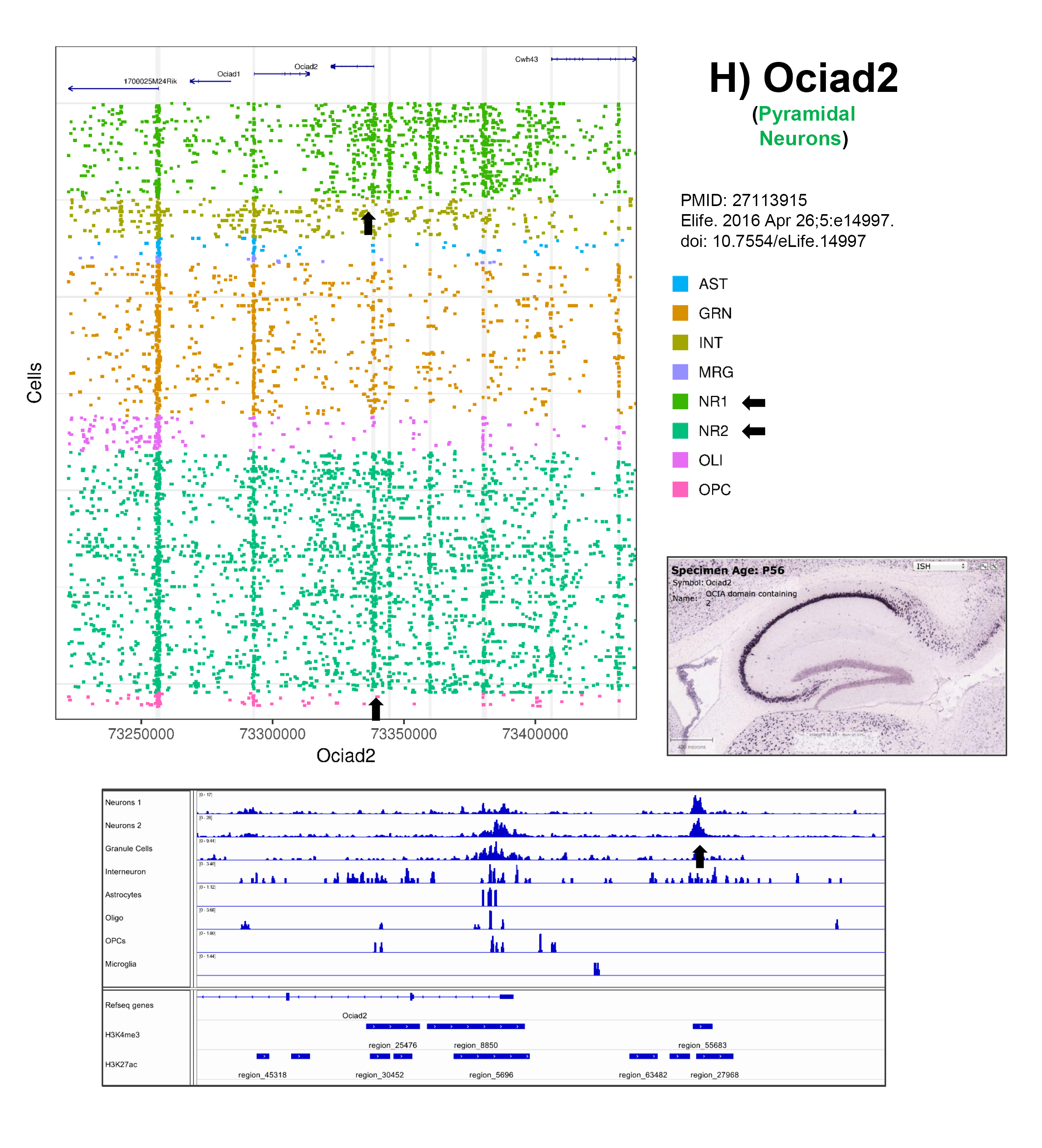
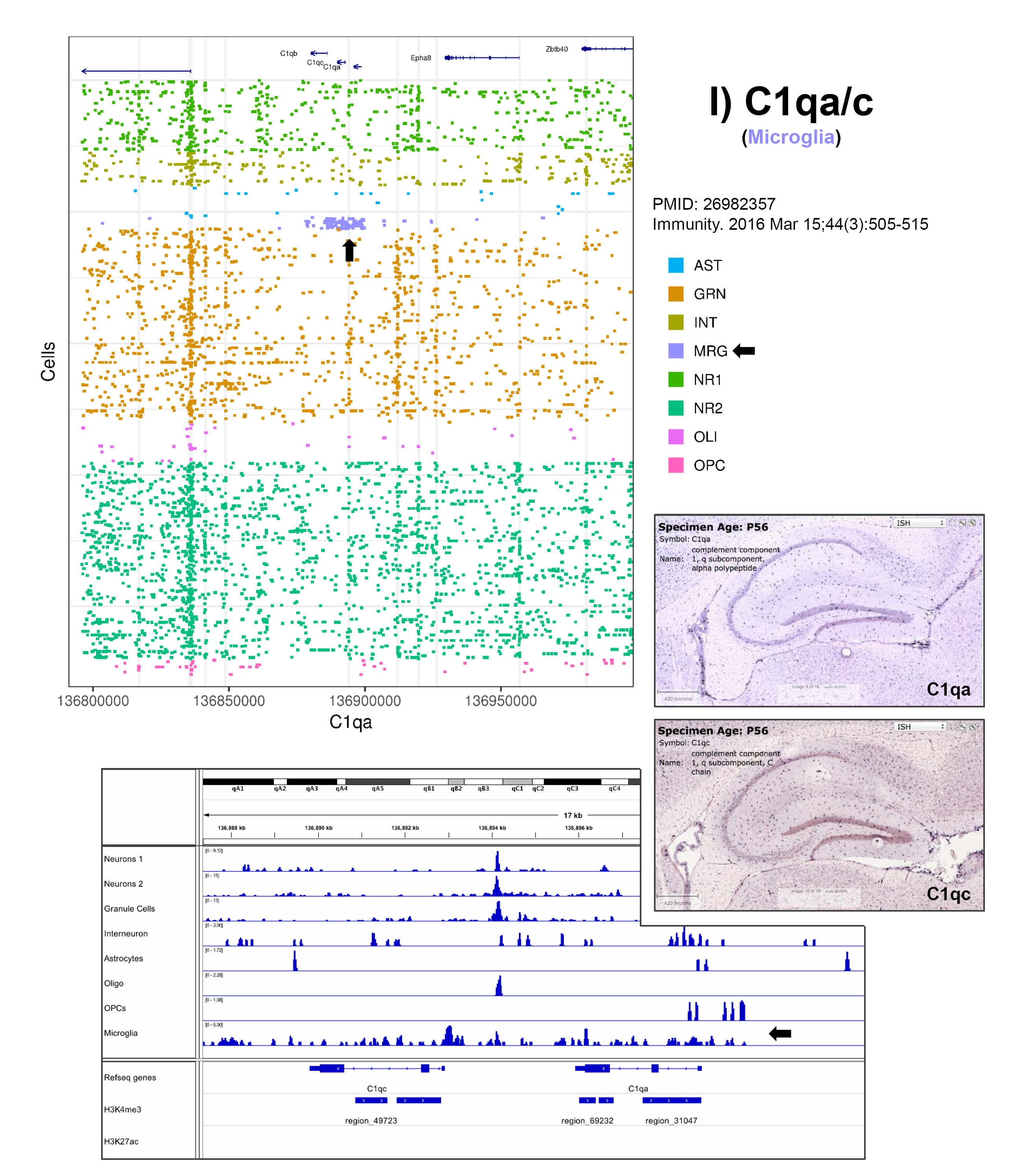
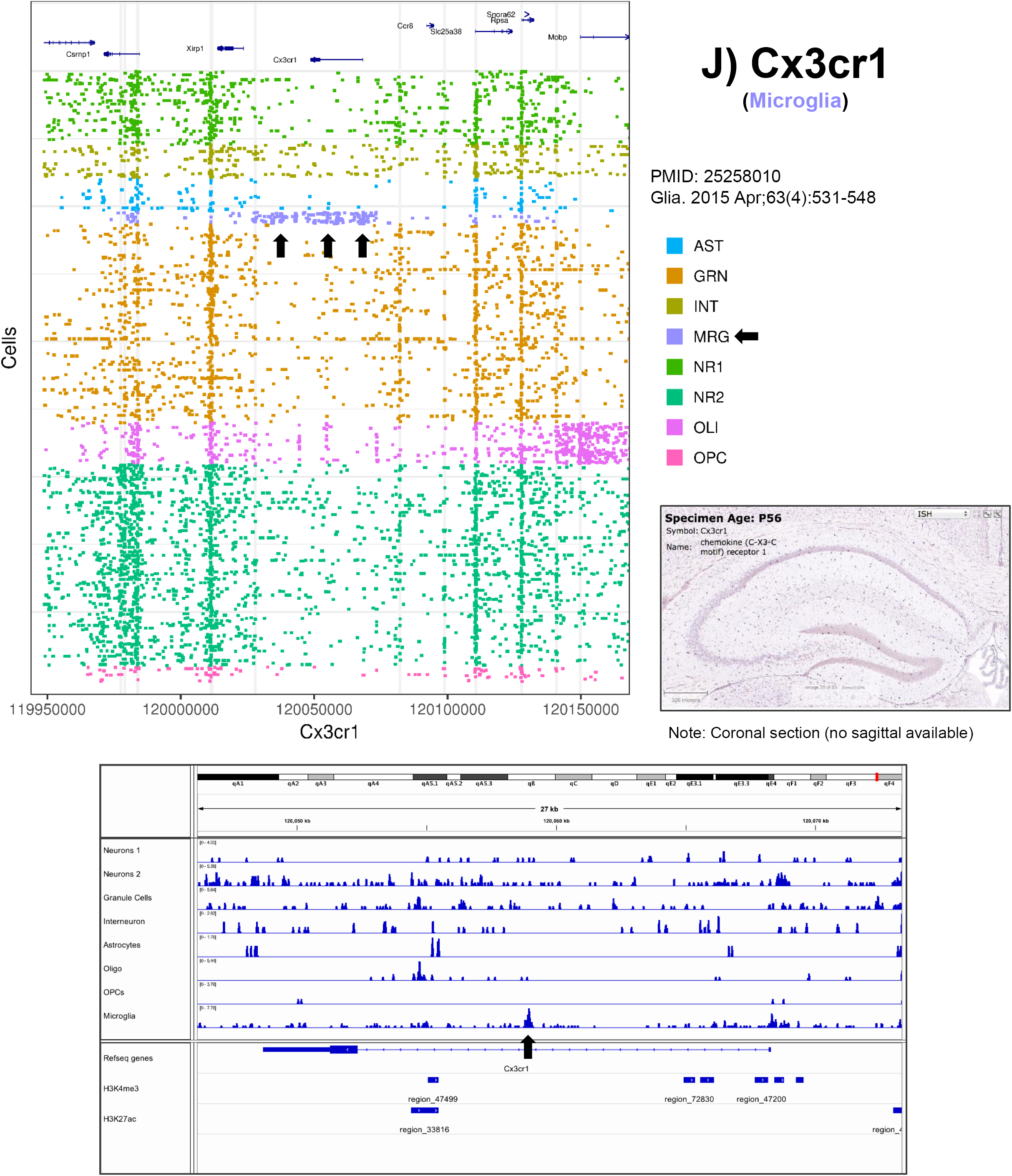
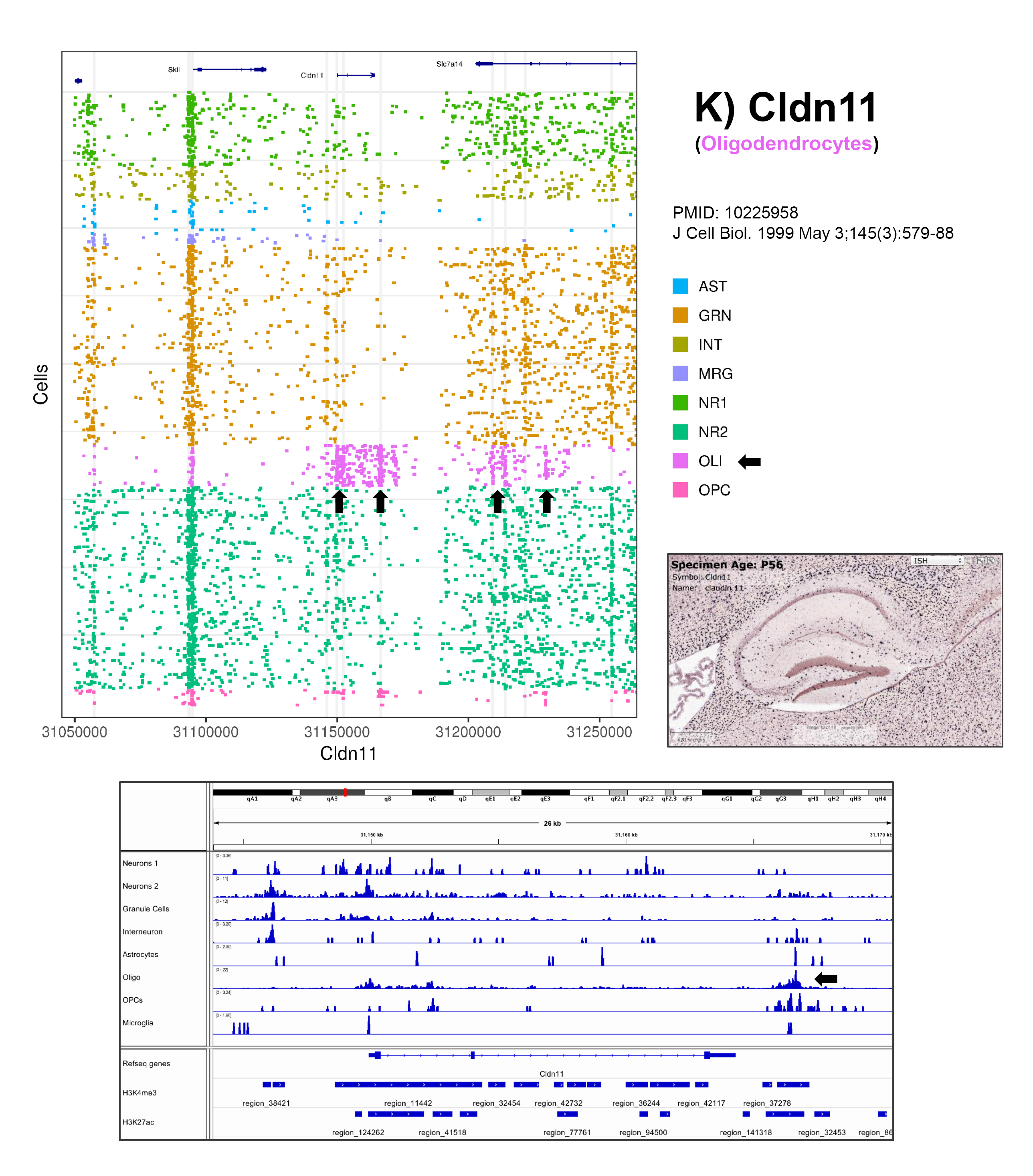
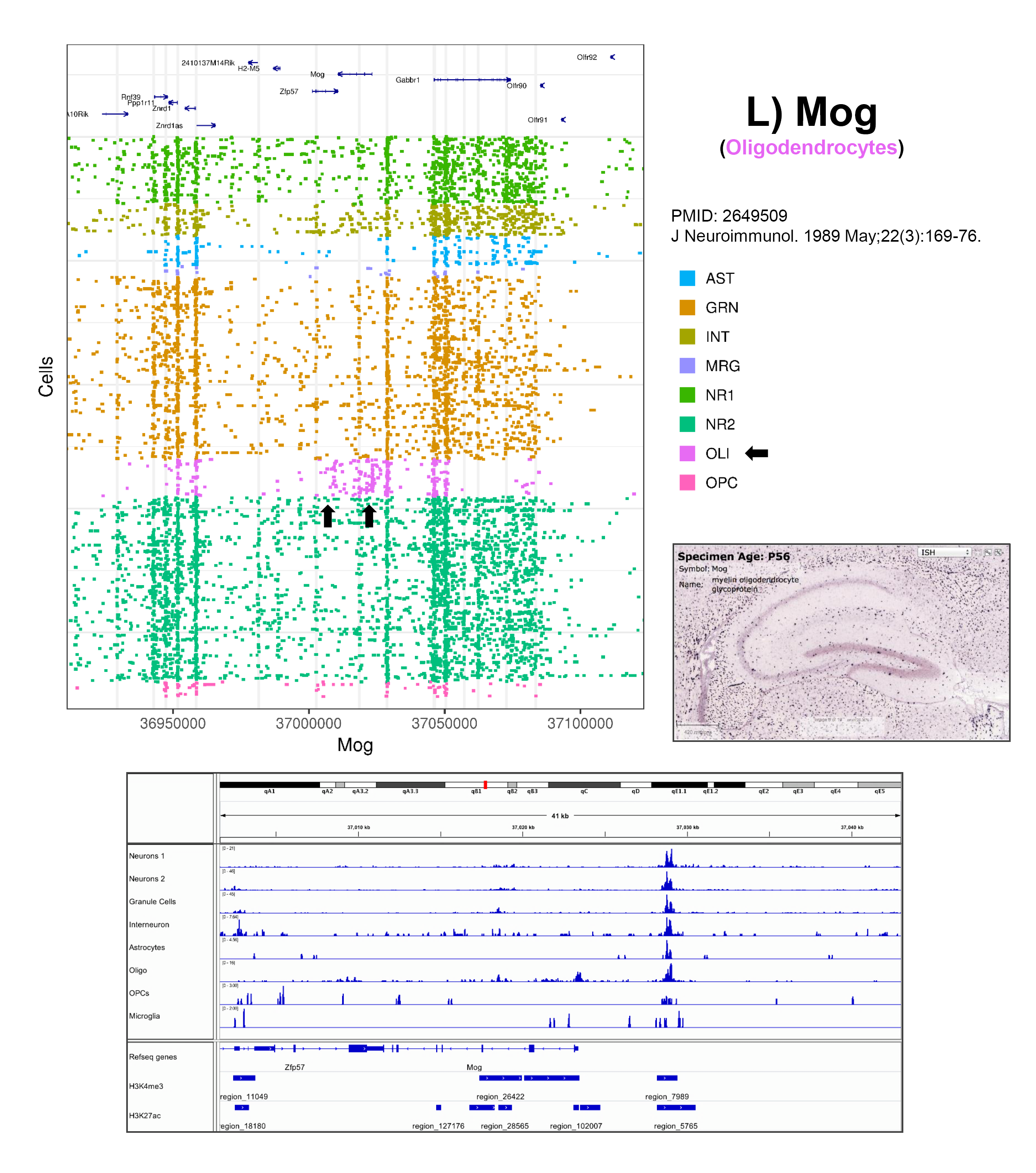
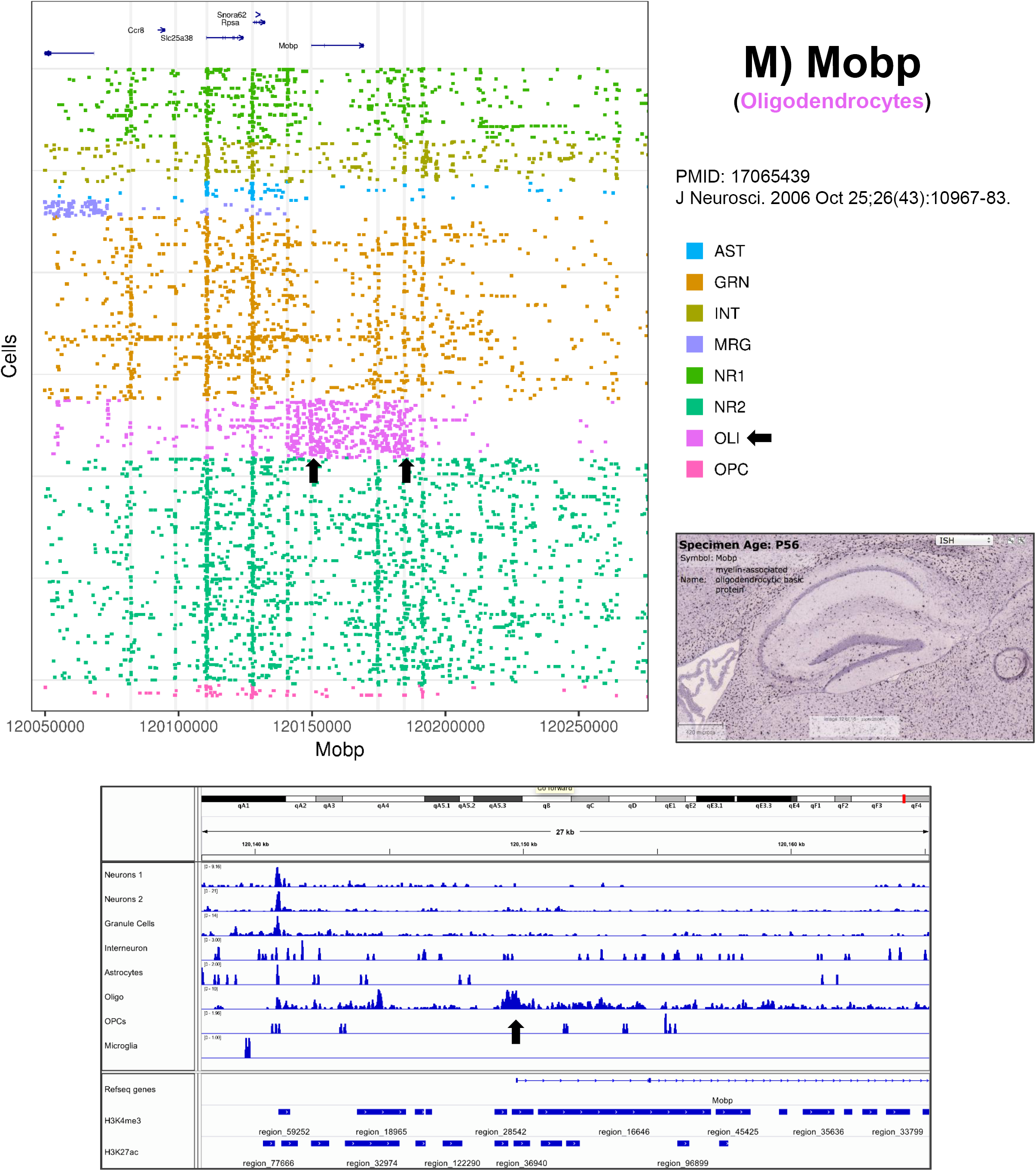
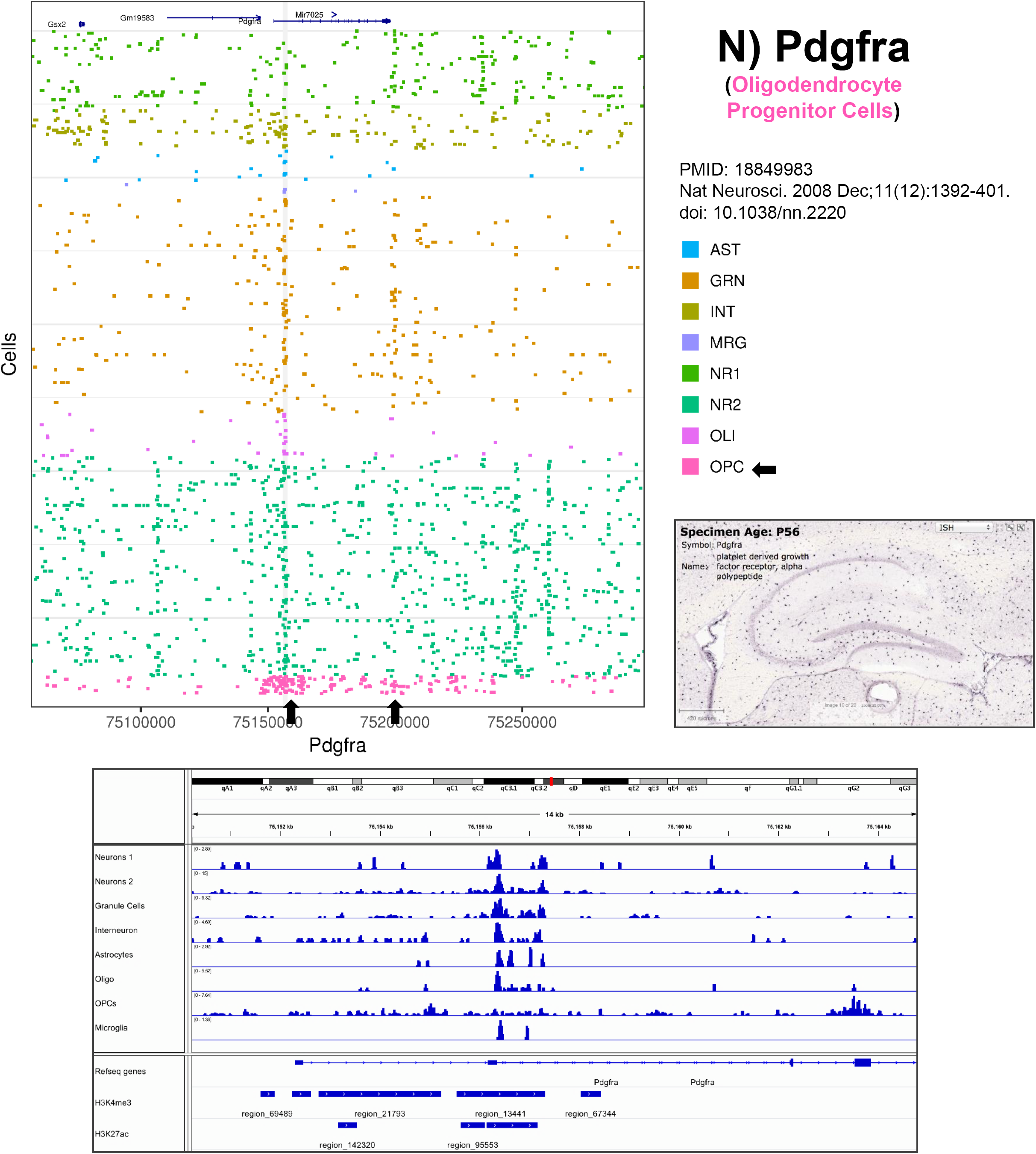
Marker Gene Dashboards. Marker gene dashboards (A-N) contain three plots and additional information. For each dashboard the gene name can be found in the upper right with the specified cell type below in parentheses. The literature reference for why the gene was selected is included below the gene title. The top left plot is a ‘read plot’ of the region around the gene (+/-100,000 bp) with genes in the region plotted at the top followed by rows for each cell with unique reads in the region colored by the cluster identity. To the right of the read plot is an immunohistochemistry image from the Allen Brain Atlas for the specified gene. Lastly, the bottom panel is a genome browser view showing the aggregated cluster sci-ATAC-seq profiles at a zoomed in region around the marker gene along with mouse hippocampus H3K4me3 and H3K27ac ChiP-seq peaks obtained from the ENCODE project. Black arrows on the read plot and genome viewer screenshot indicate cluster-specific signal and the corresponding cluster in the legend of the read plot.

**Supplementary Figure 2:**
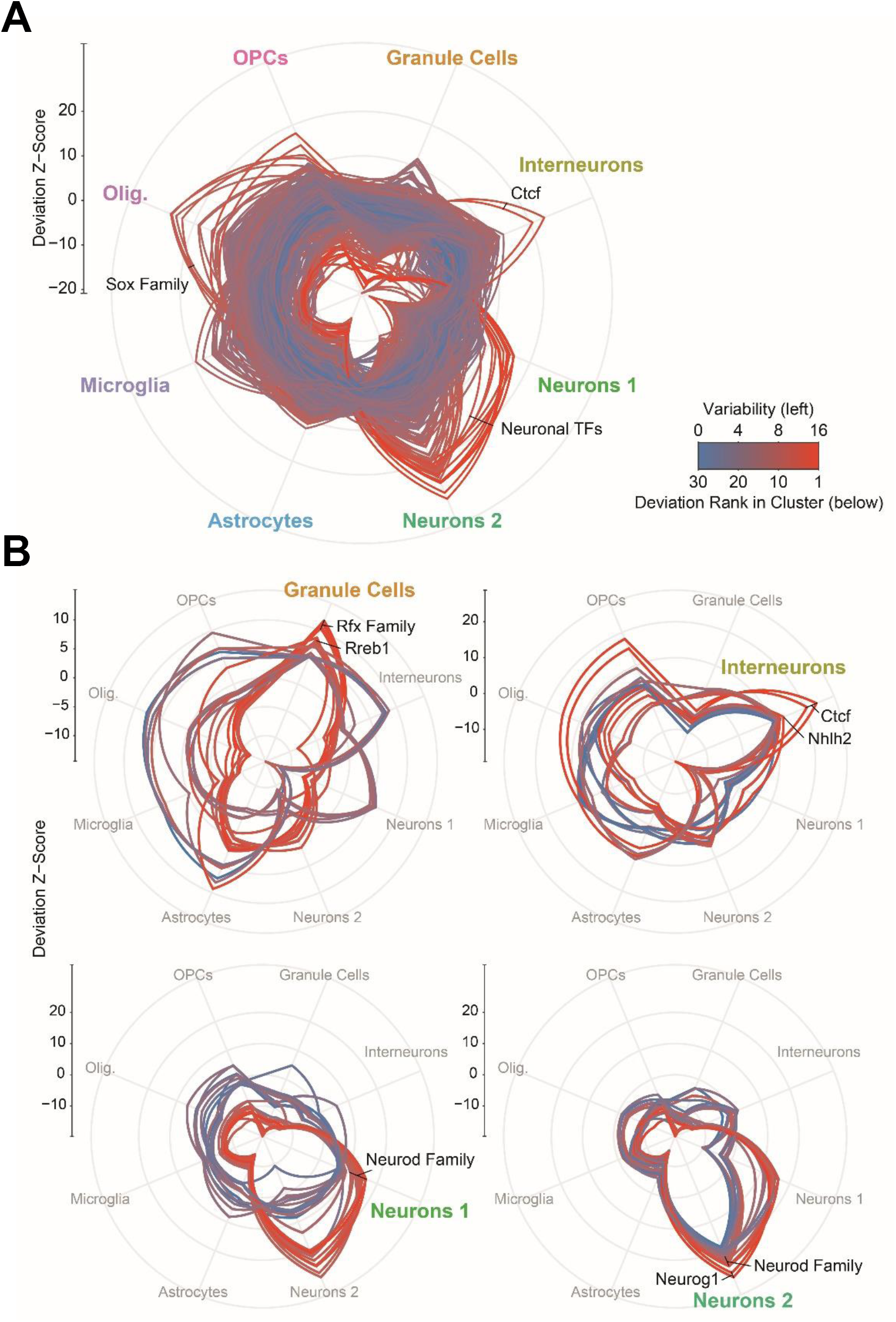
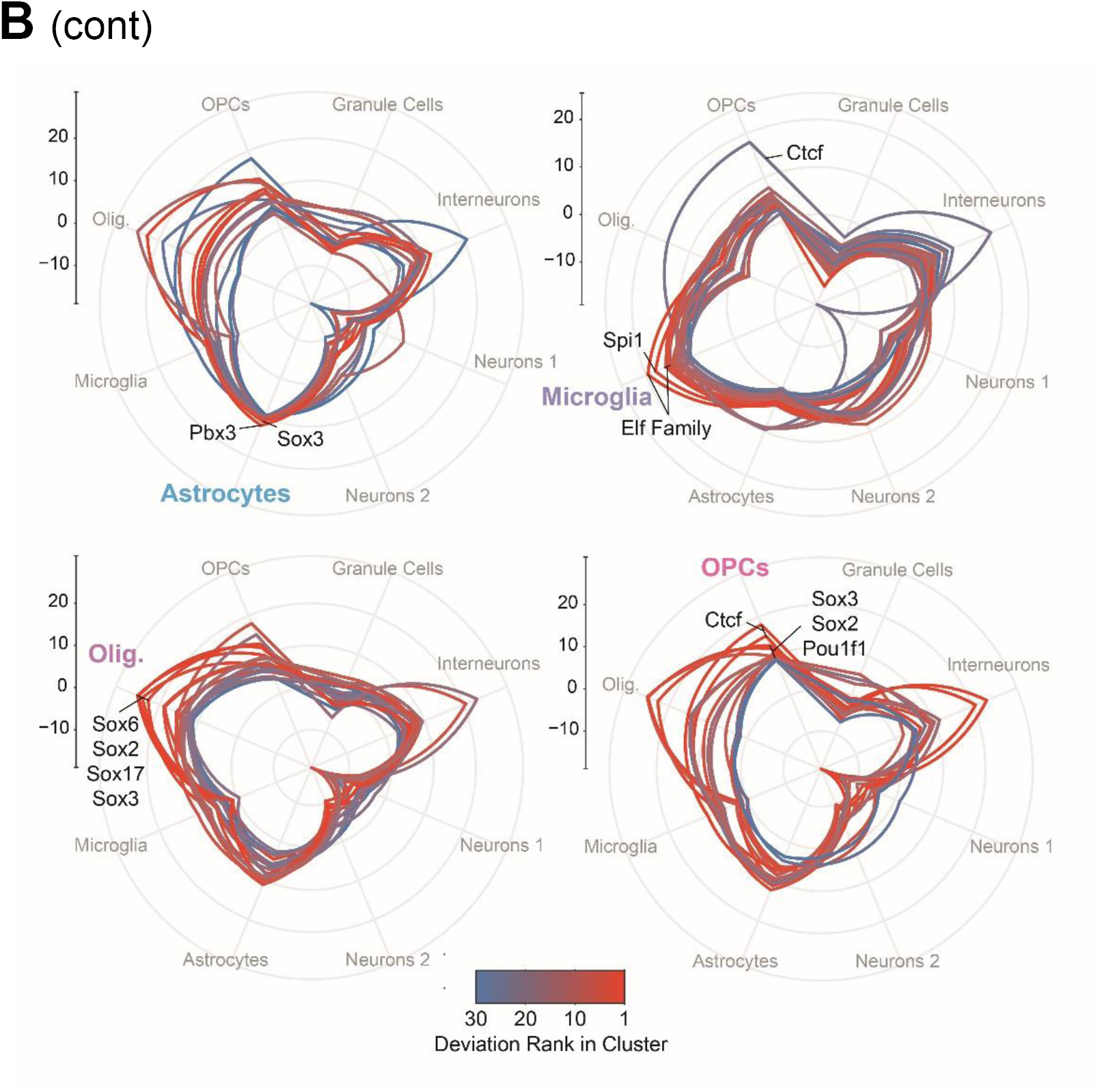
Global motif profiles for cell types. A) *ChromVAR* mean deviation scores (y-axis) for each motif for each cell type. Color indicates the variability score for the motif as reported by *ChromVAR*. B) Top 30 motifs with the highest deviation z-scores as shown in A. Color indicates the ranking within the cluster. Polygon plots were utilized to confer similar shapes of top motif accessibility between clusters.

**Supplementary Figure 3:**
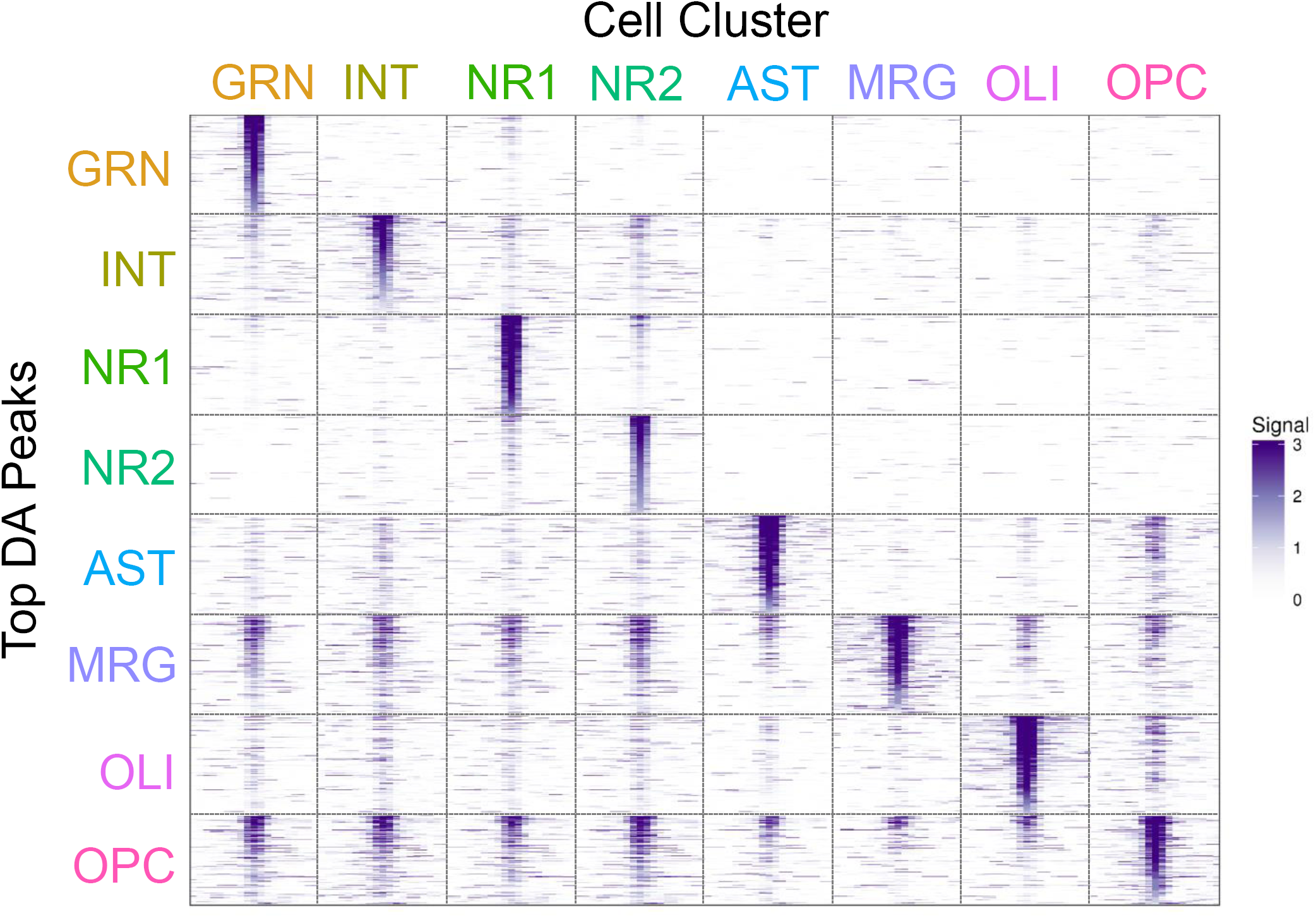
DA Peak ATAC-seq Signal. ATAC-seq signal for the top 1,000 differentially accessible peaks for each cell type are shown as in Fig. 2 but sorted according to peaks with the top signal versus significance of differential accessibility.

**Supplementary Figure 4:**
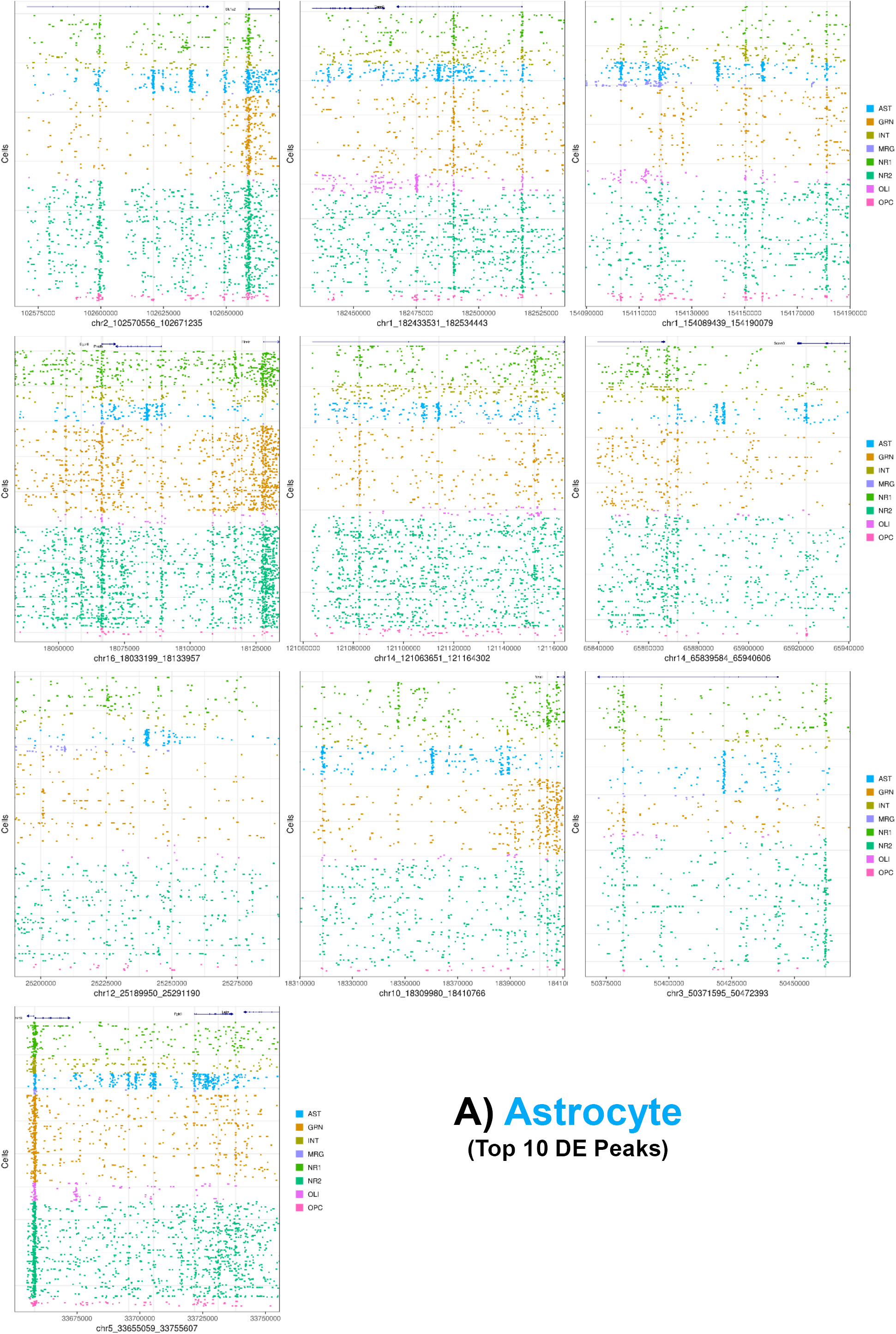
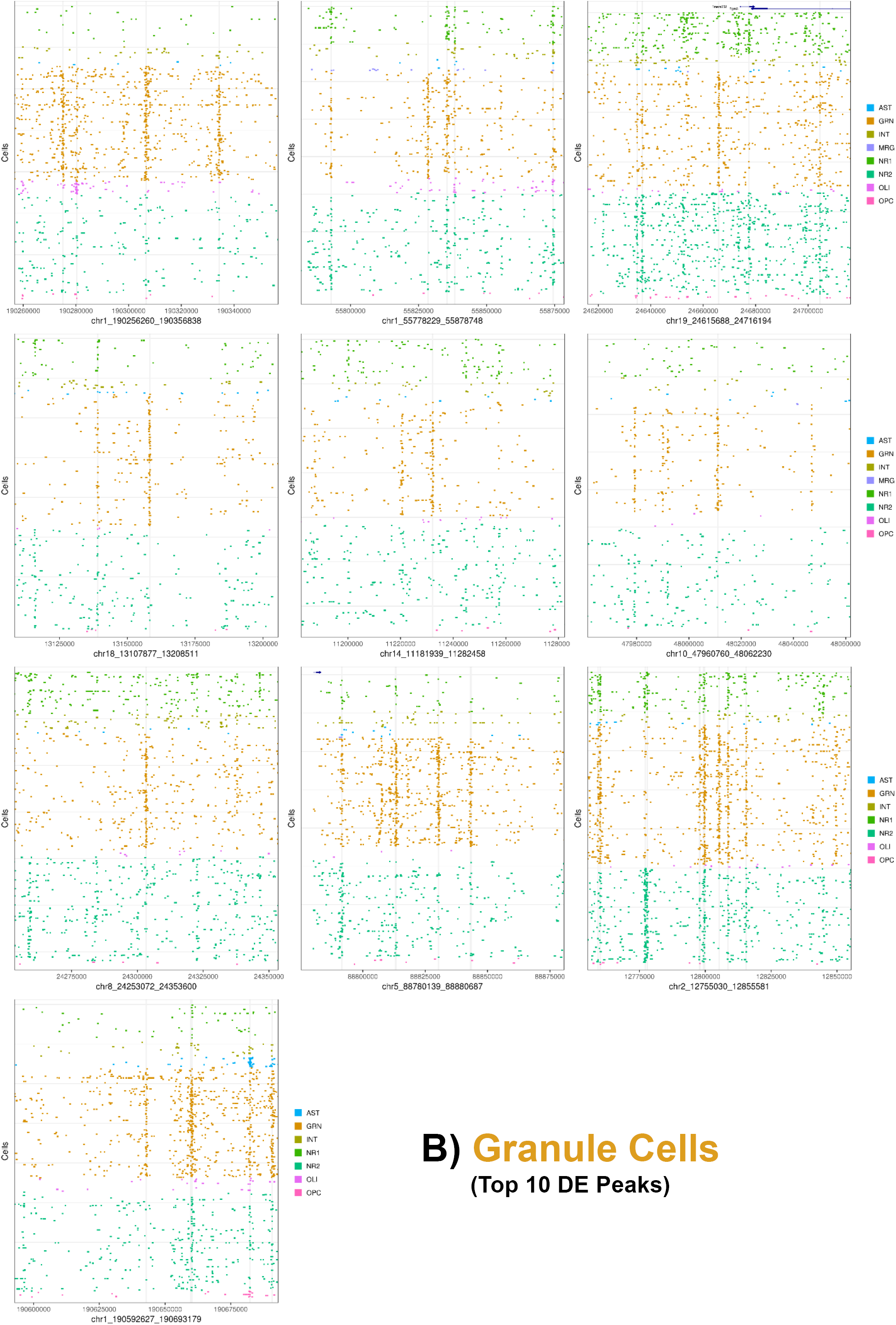
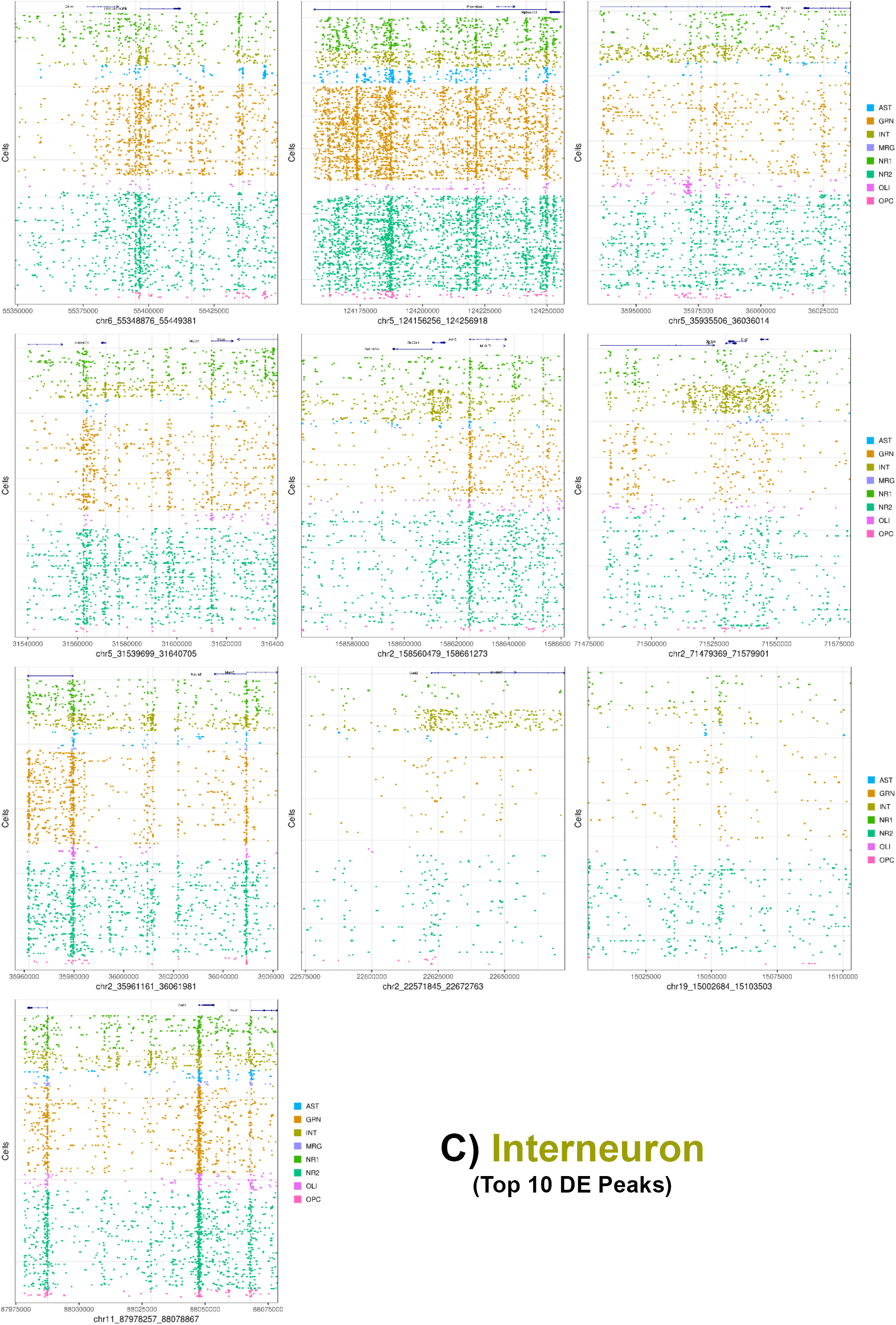
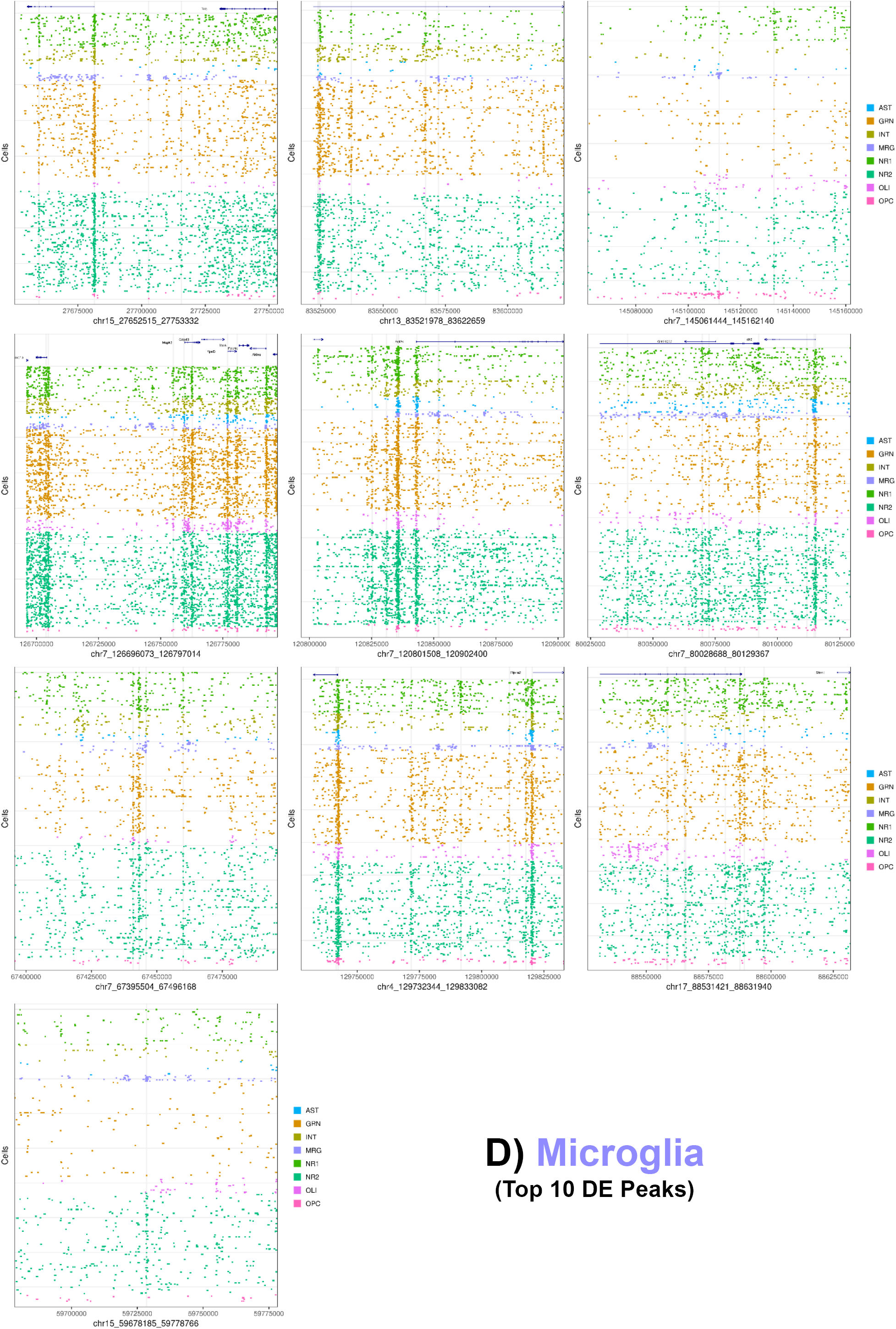
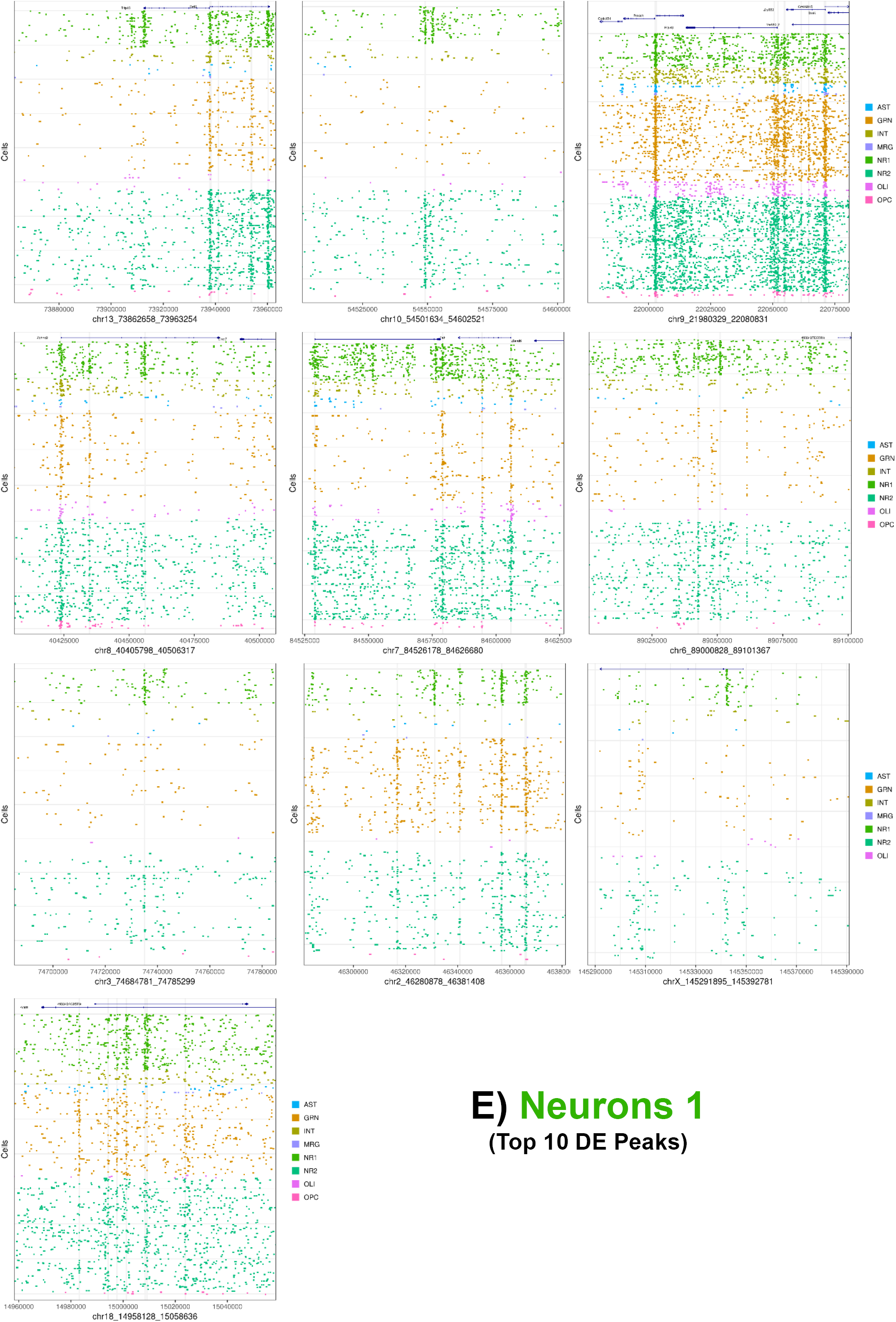
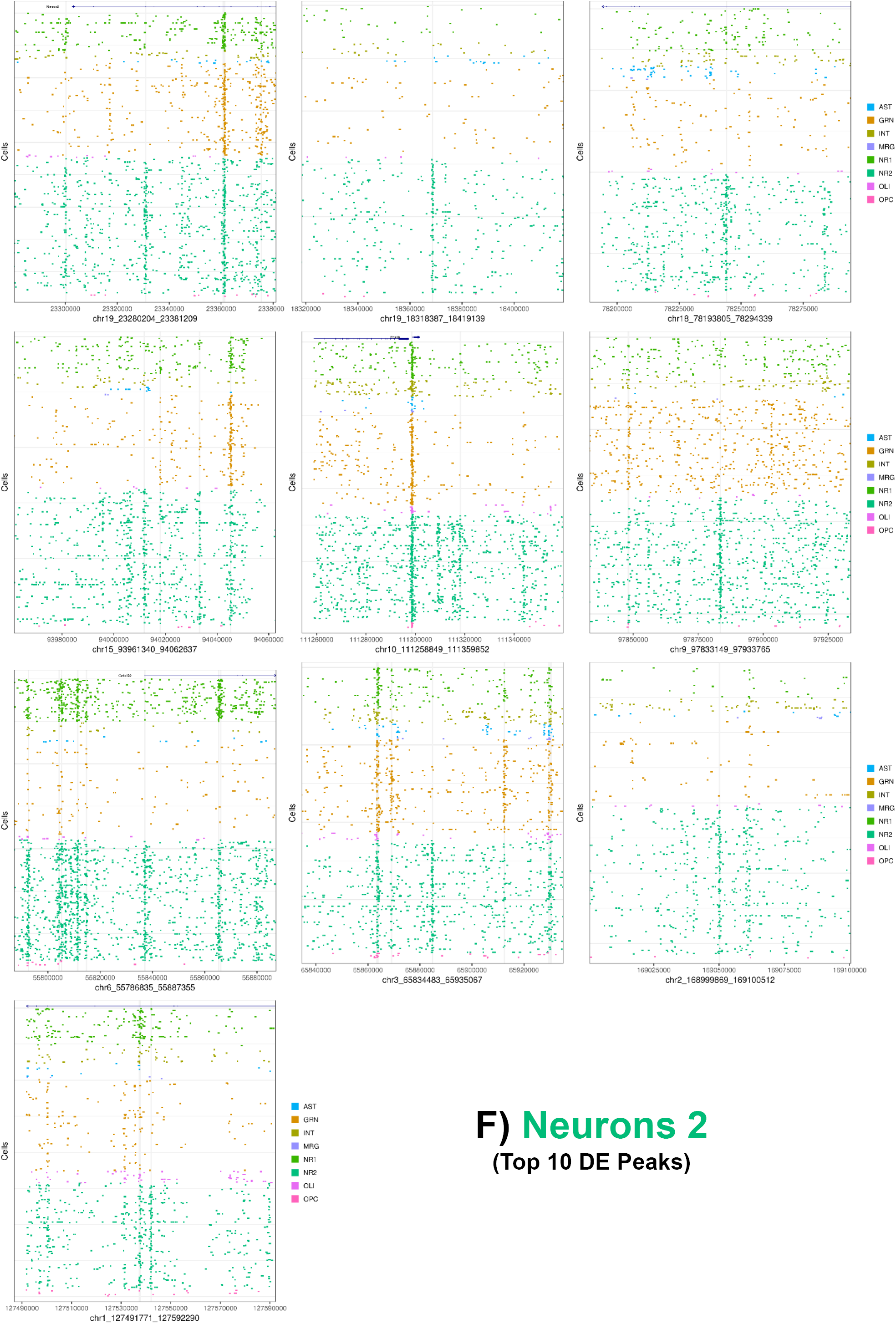
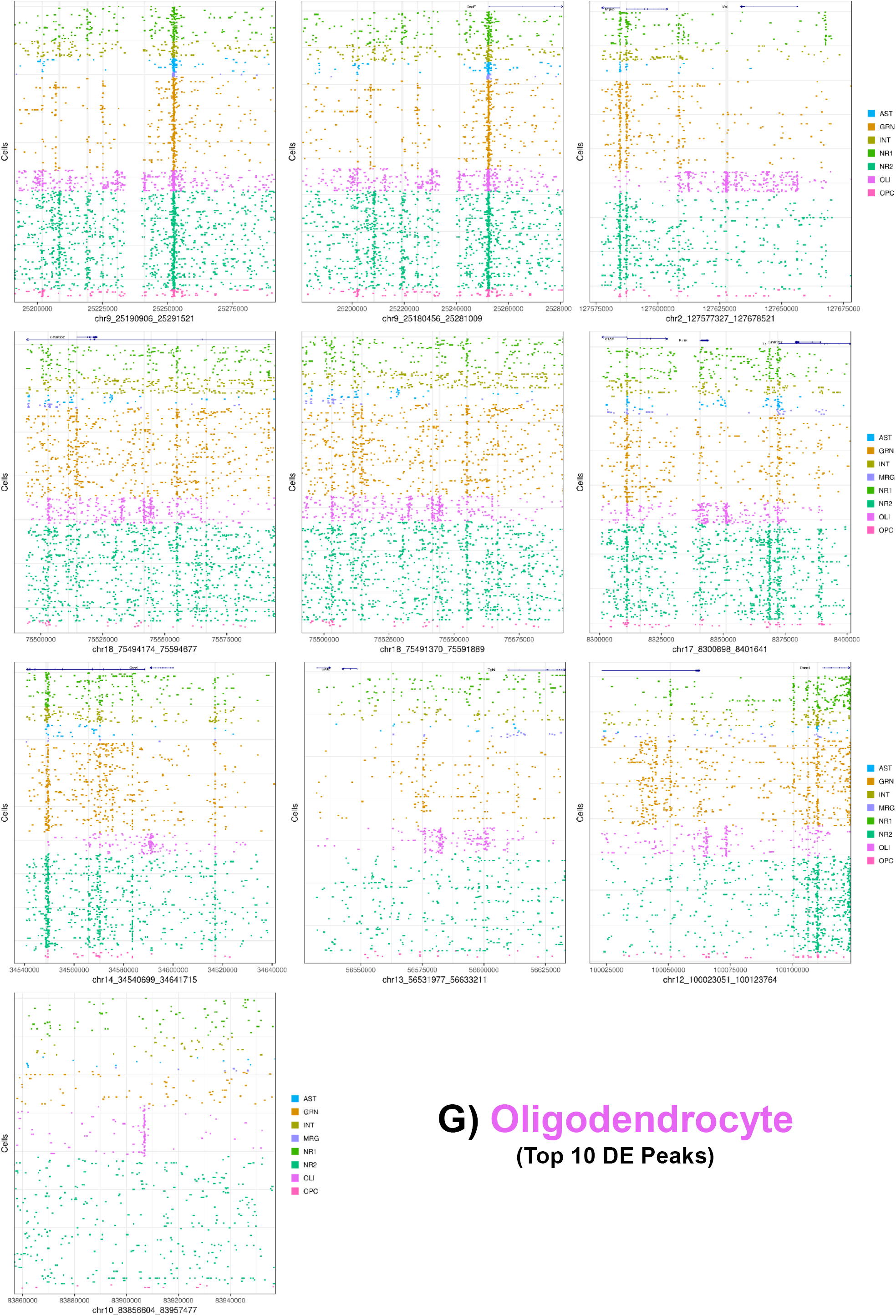
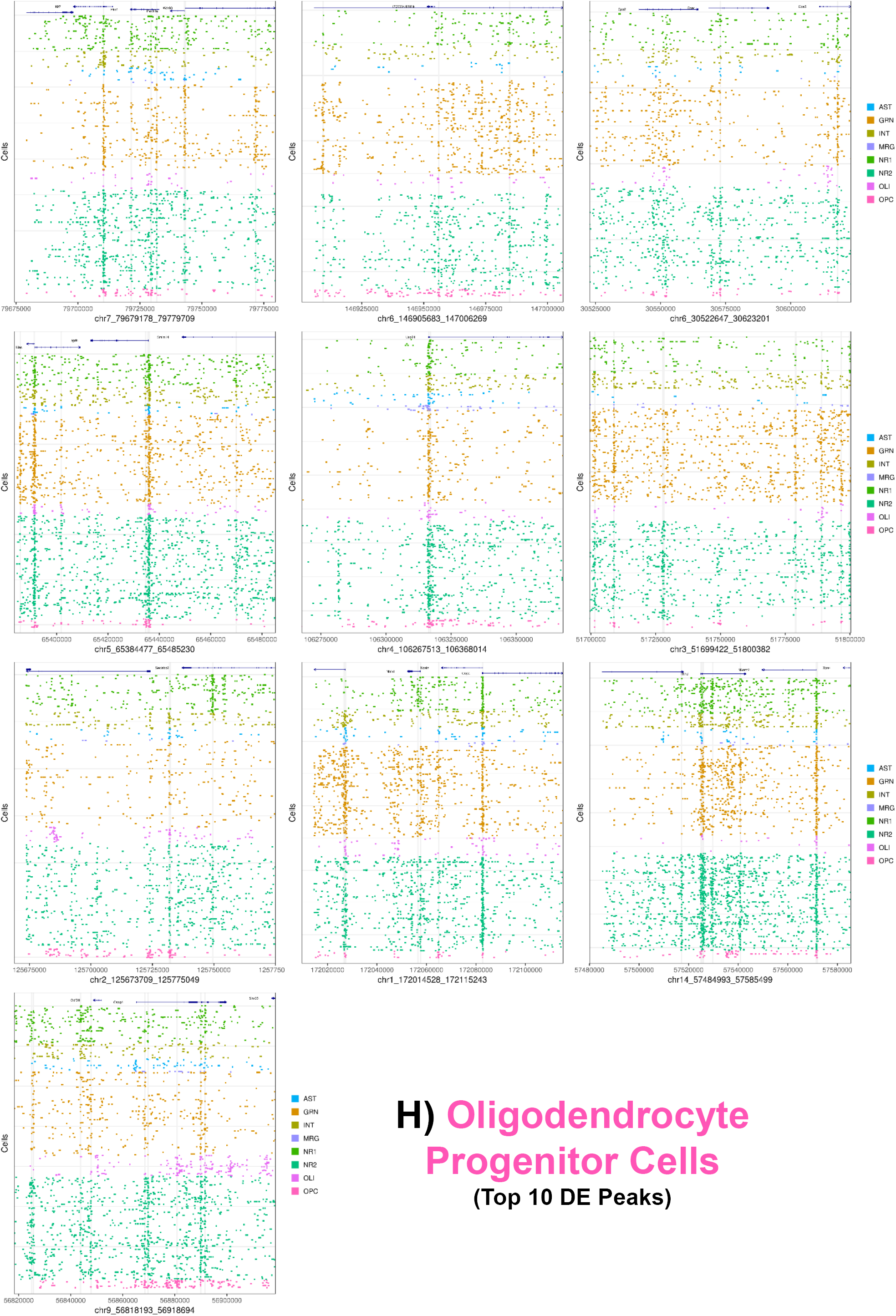
Differentially Accessible Peaks. The top 10 differentially accessible (Supplementary Table 2, see Methods) peaks corresponding to each cluster (A-H) are plotted along with the sci-ATAC-seq reads present within the region +/-50,000 basepairs of the identified DA peak (centered).

**Supplementary Figure 5:**
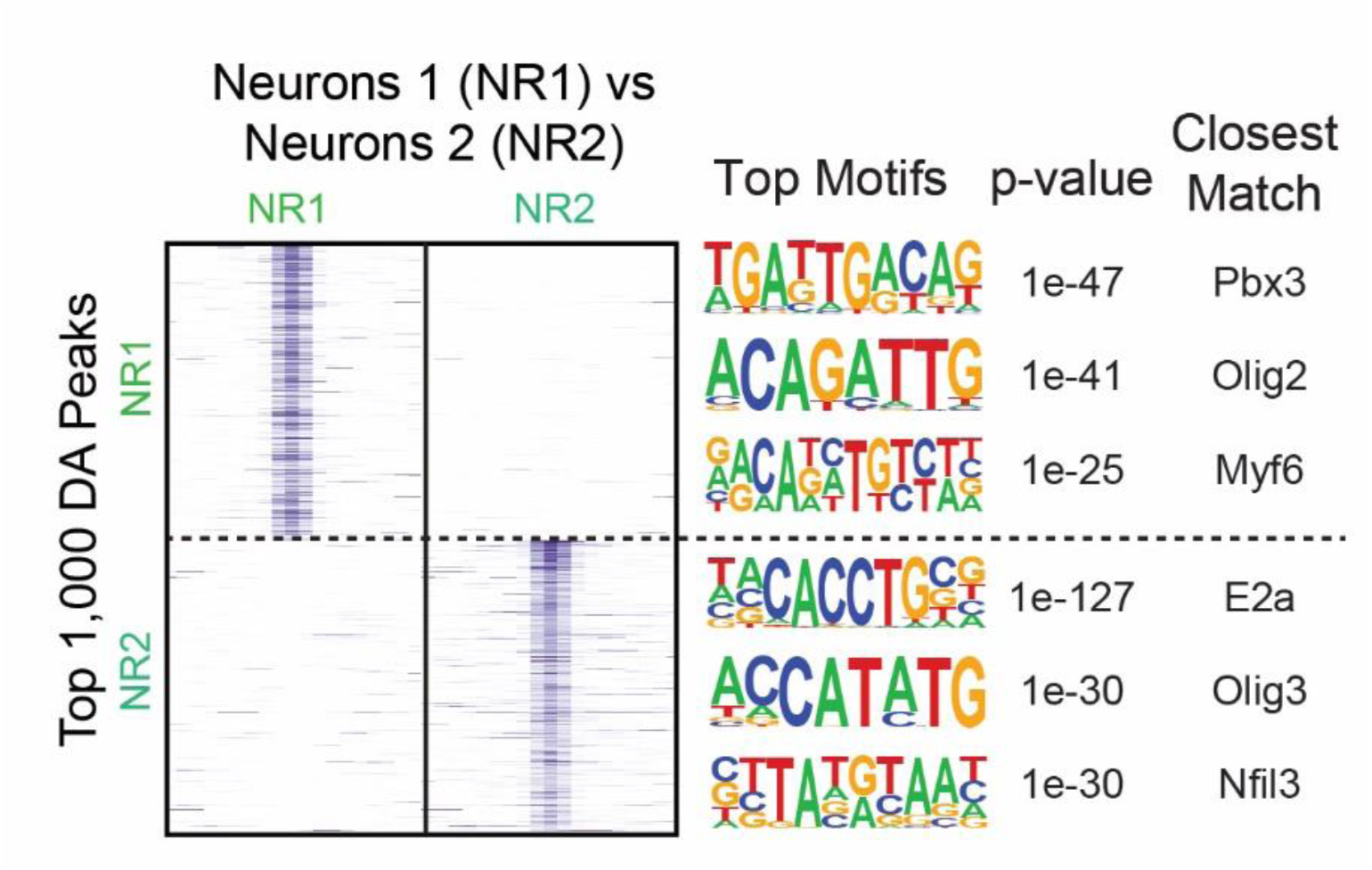
Top DA peaks between NR1 and NR2 excitatory neuron cell types. ATAC-seq signal for the top 1,000 DA peaks is plotted for each cell type along with the top three motifs associated with each peak set and their corresponding p-value and closest matching motif.

**Supplementary Figure 6:**
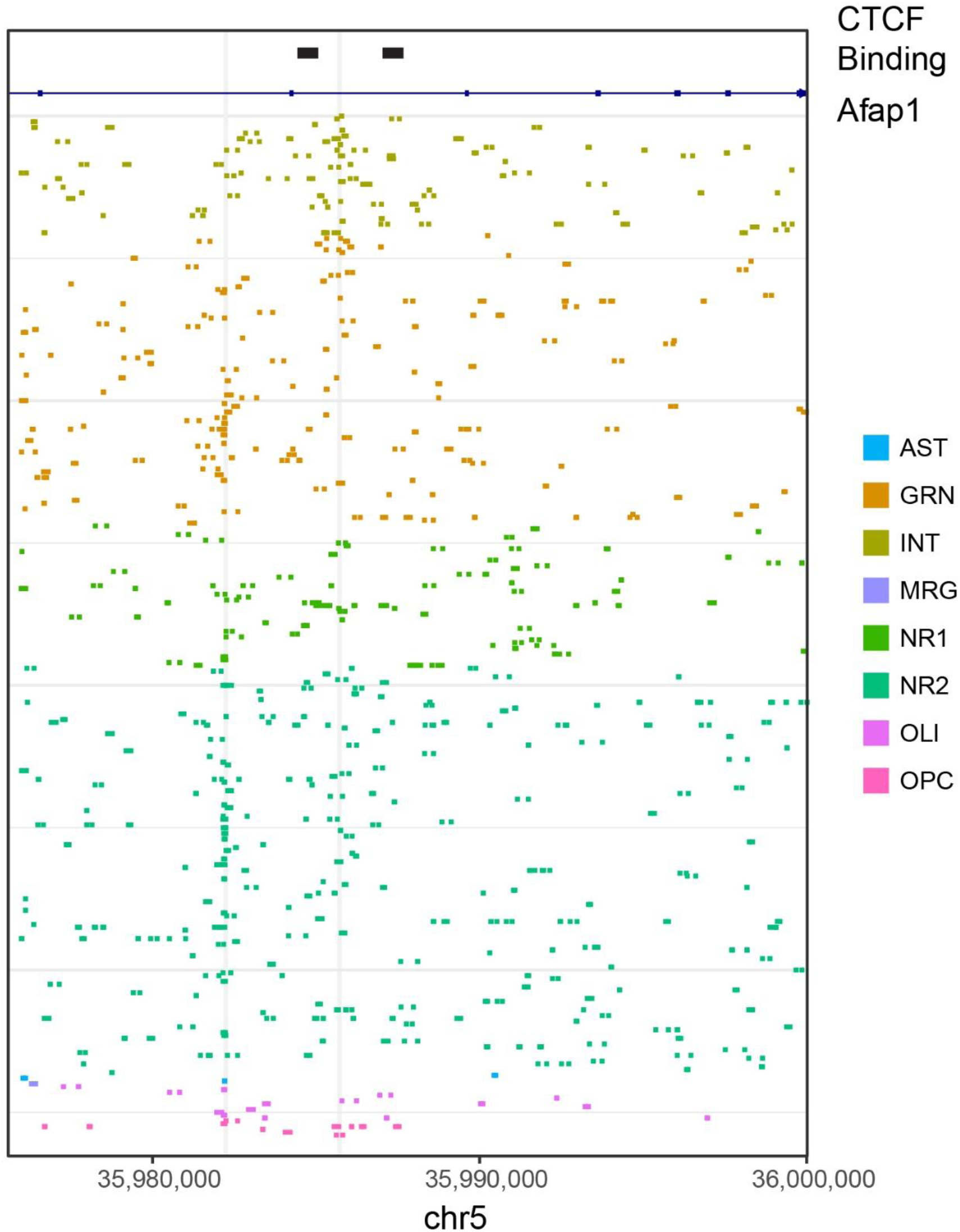
Afap1 locus. Read plot generated by scitools showing a peak highly enriched in interneurons that is flanked by CTCF ChIP-seq peaks.

**Supplementary Figure 7:**
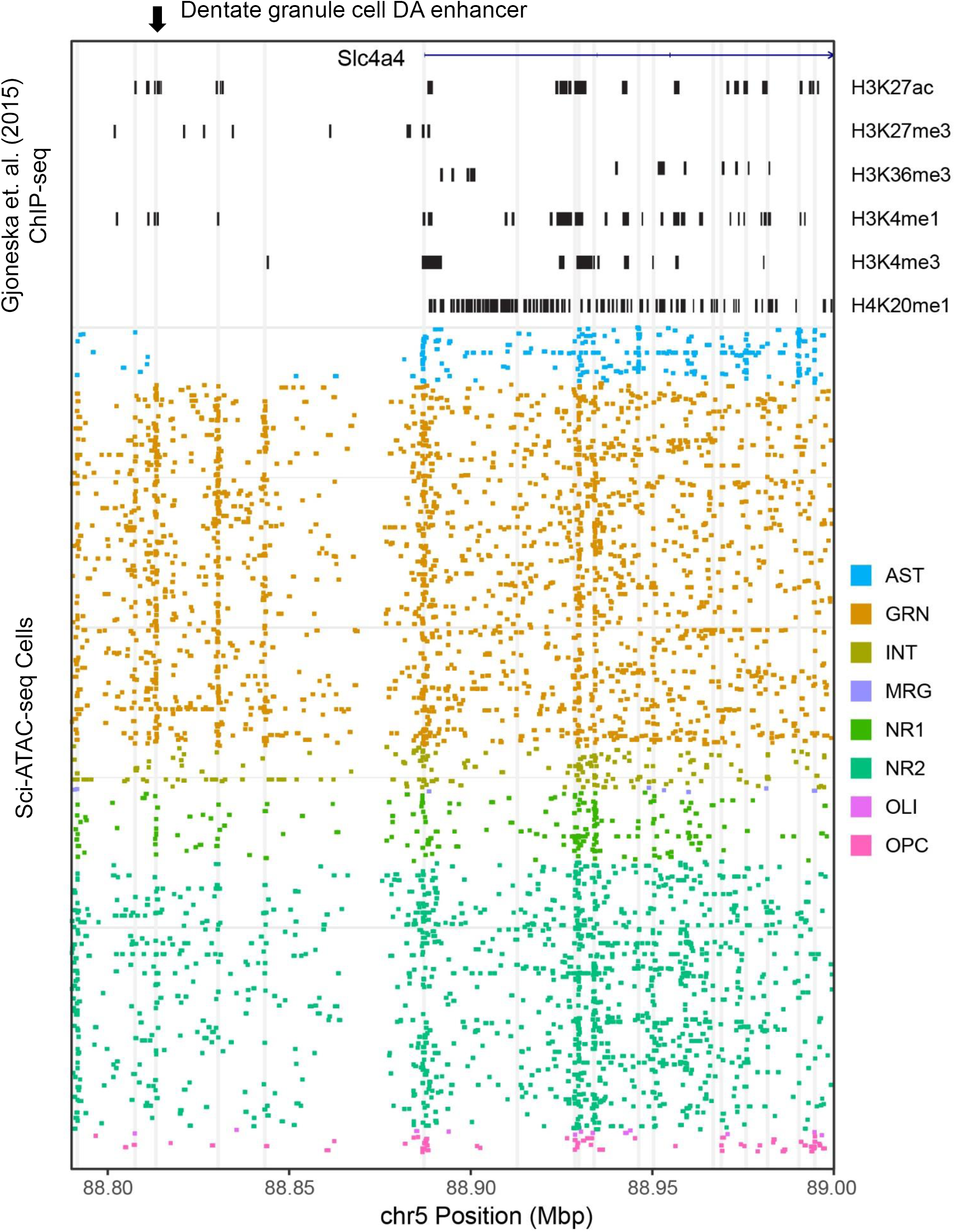
Slc4a4 locus. Read plot generated by scitools showing the putative cell-type-specific enhancer of *Slc4a4* that is differentially accessible in the dentate granule cell population. ChIP-seq peaks from Gjoneska et. al. (2015) are shown below the gene track in black.

**Supplementary Figure 8:**
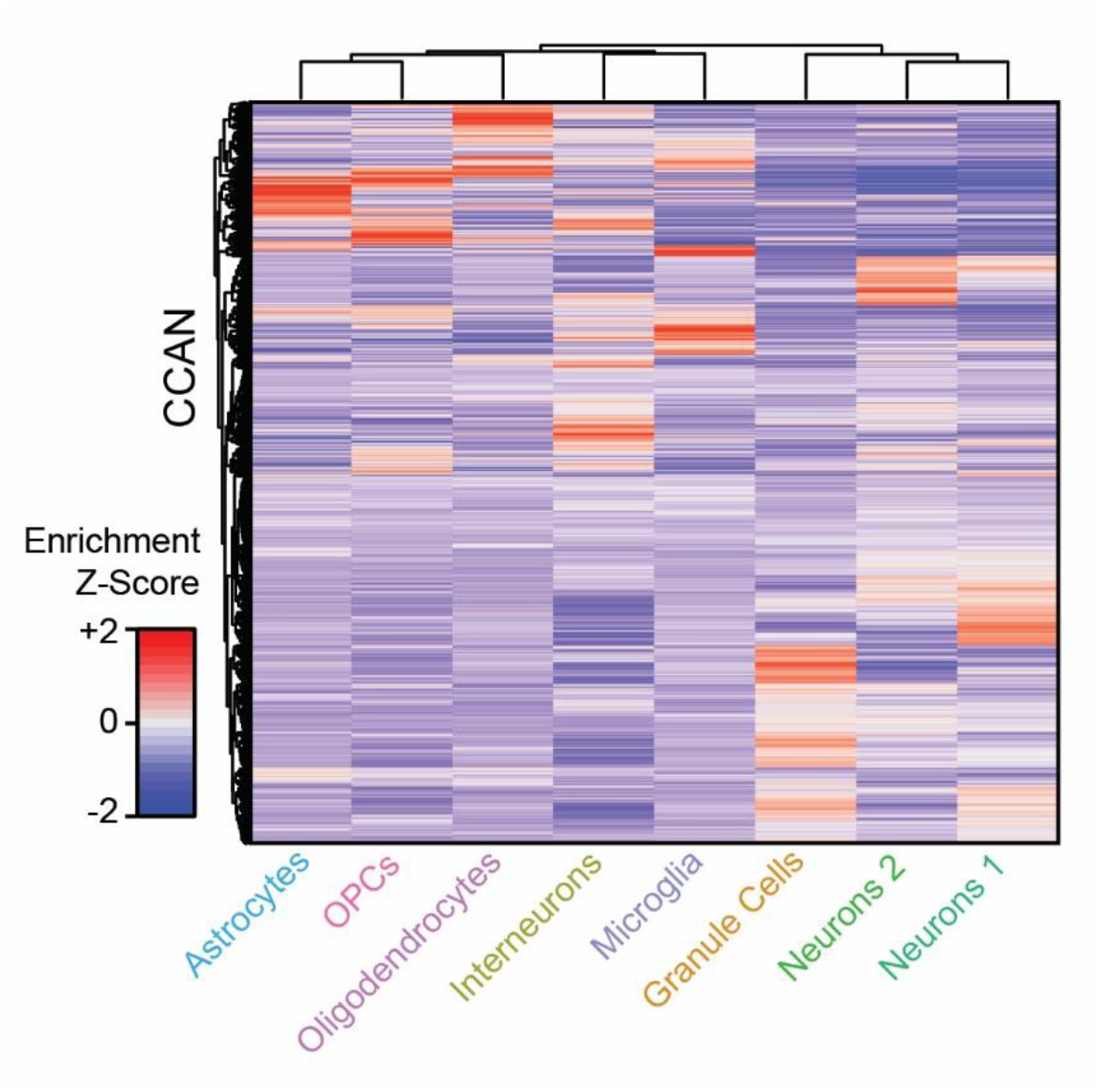
Hierarchical clustering of CCANs by cell type enrichment. Hierarchical clustering of enrichment z-scores for peaks contained with in each CCAN with respect to cell type.

**Supplementary Figure 9:**
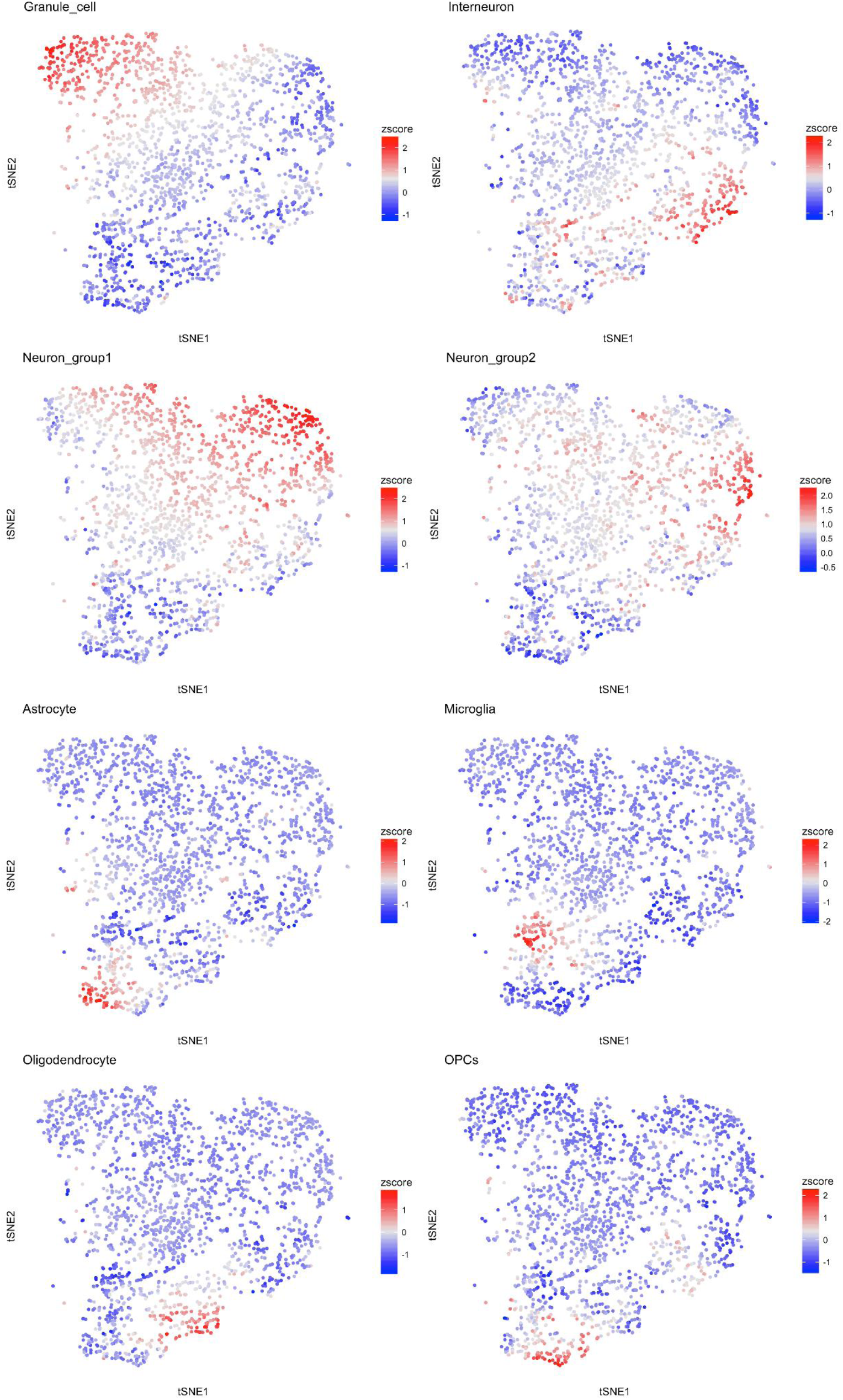
CCAN cell type enrichments on tSNE projections. Enrichment for CCAN’s within each cell type is plotted as in Fig. 3, but for each of the eight major clusters.

**Supplementary Figure 10:**
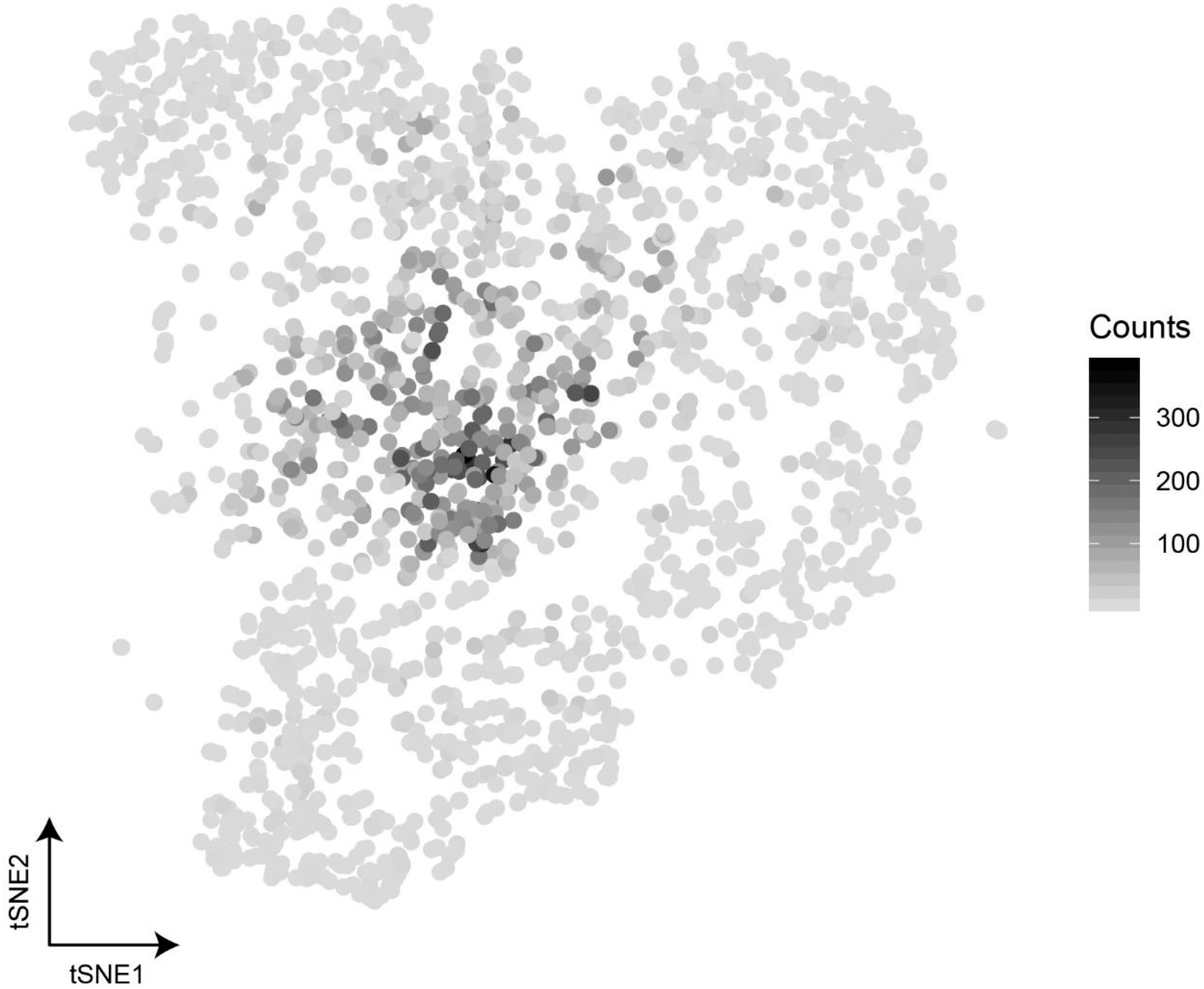
CCAN peak counts. tSNE of CCANs by cell type specificity (as in Figure 3), but shaded by peak membership count. CCANs with greater numbers group towards the center and are less-cell type specific overall. This is likely due to modular CCANs where certain portions of CCANs are cell-type-specific that link up with other, more universally established CCANs, as in the Prox1 example.

**Supplementary Figure 11:**
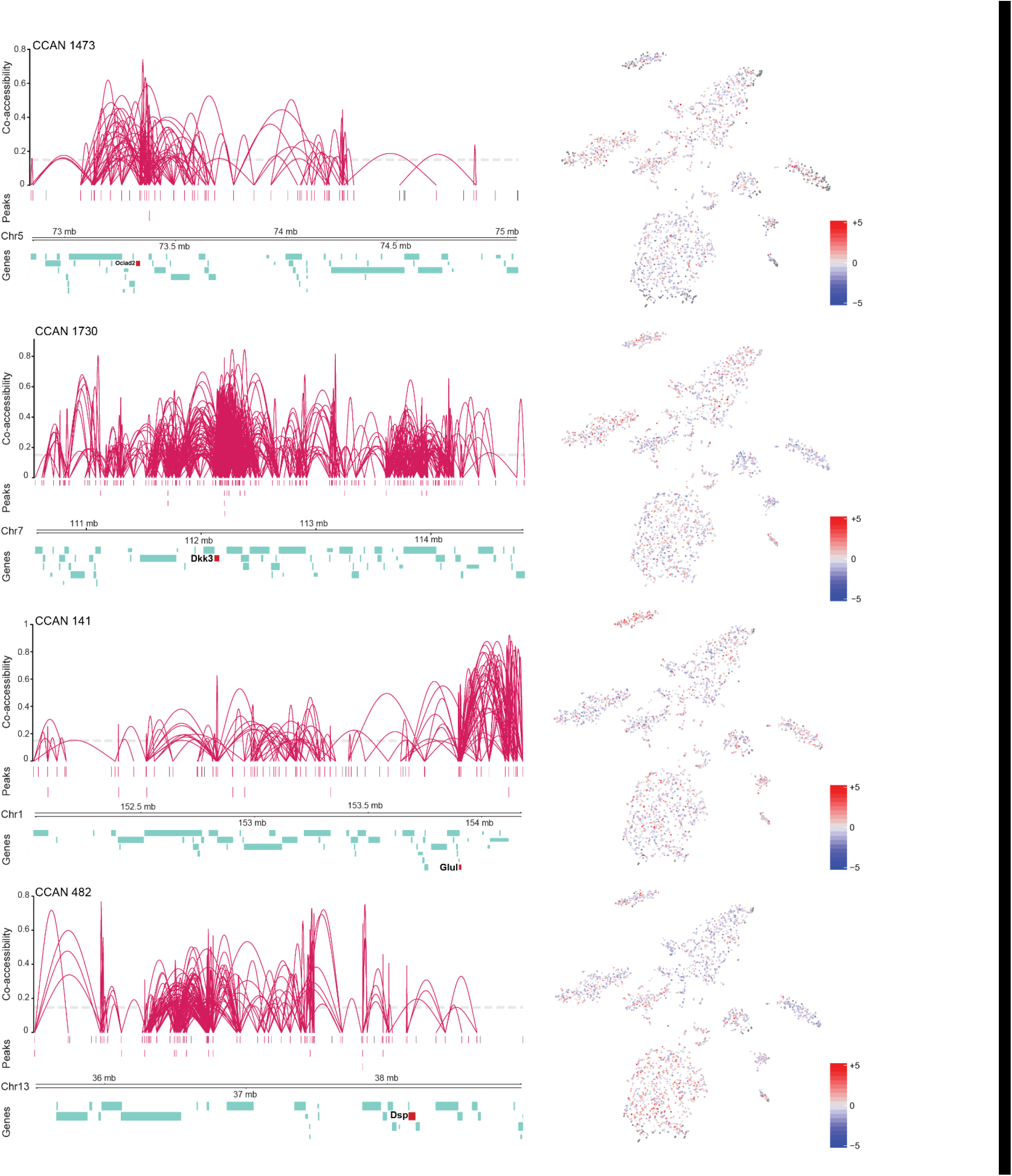
Additional cell-type-specific CCANs. Left: plots of the CCANs near cell-type-specific marker genes. Right: Accessibility enrichment across cells.

**Supplementary Figure 12:**
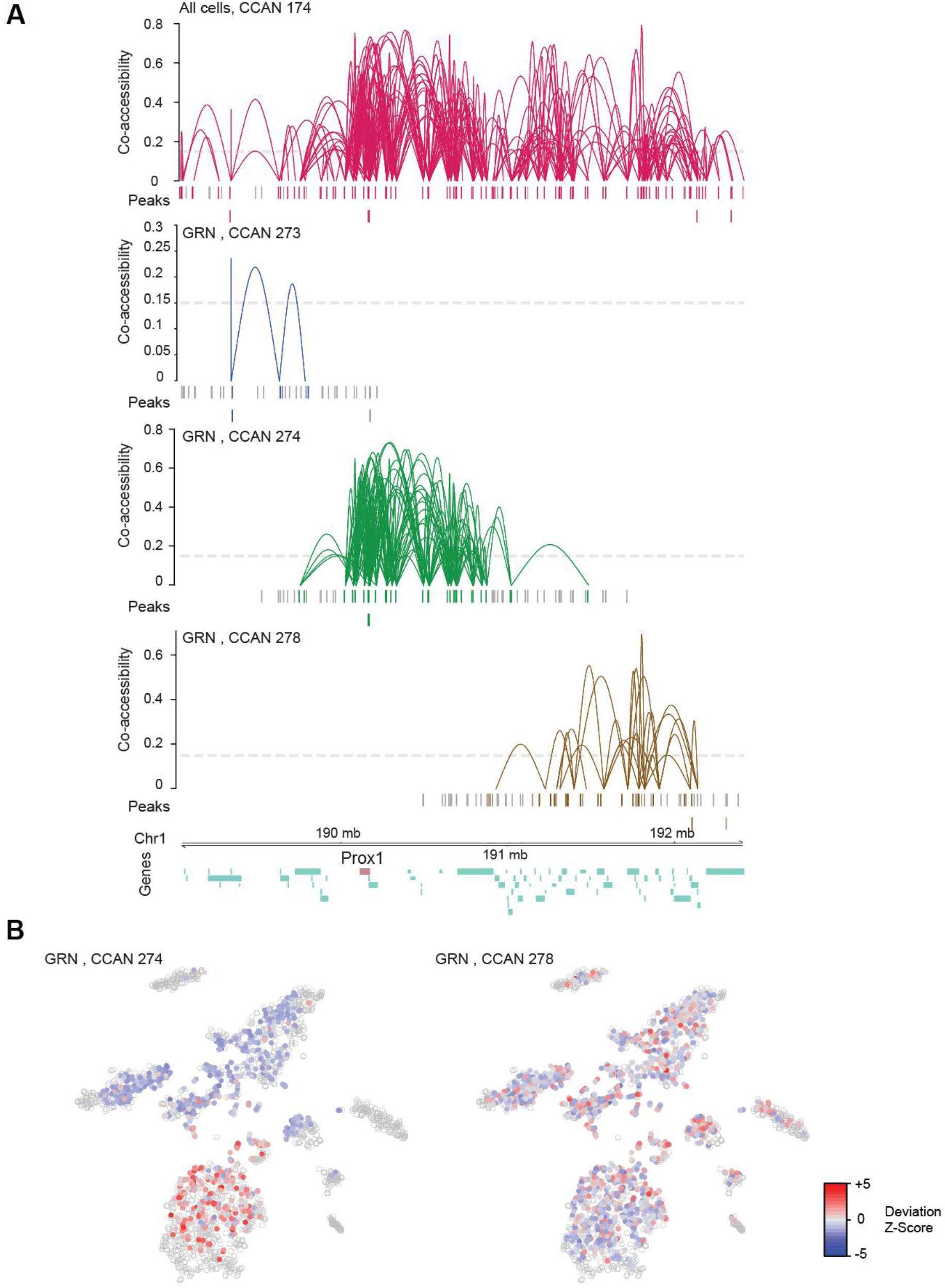
Dissection of CCAN 174 (Prox1) A) CCAN 174 as determined by Cicero performed on all cell types (top). The CCAN is large and likely contains links that are not cell-type specific. When Cicero is carried out only on the dentate granule cell population, the larger CCAN is split into three distinct networks. B) The cell-type specificity for the granule-specific CCAN (274) centered on Prox1 is more accessible than the adjacent CCAN that does not exhibit cell type specificity (278).

**Supplementary Figure 13:**
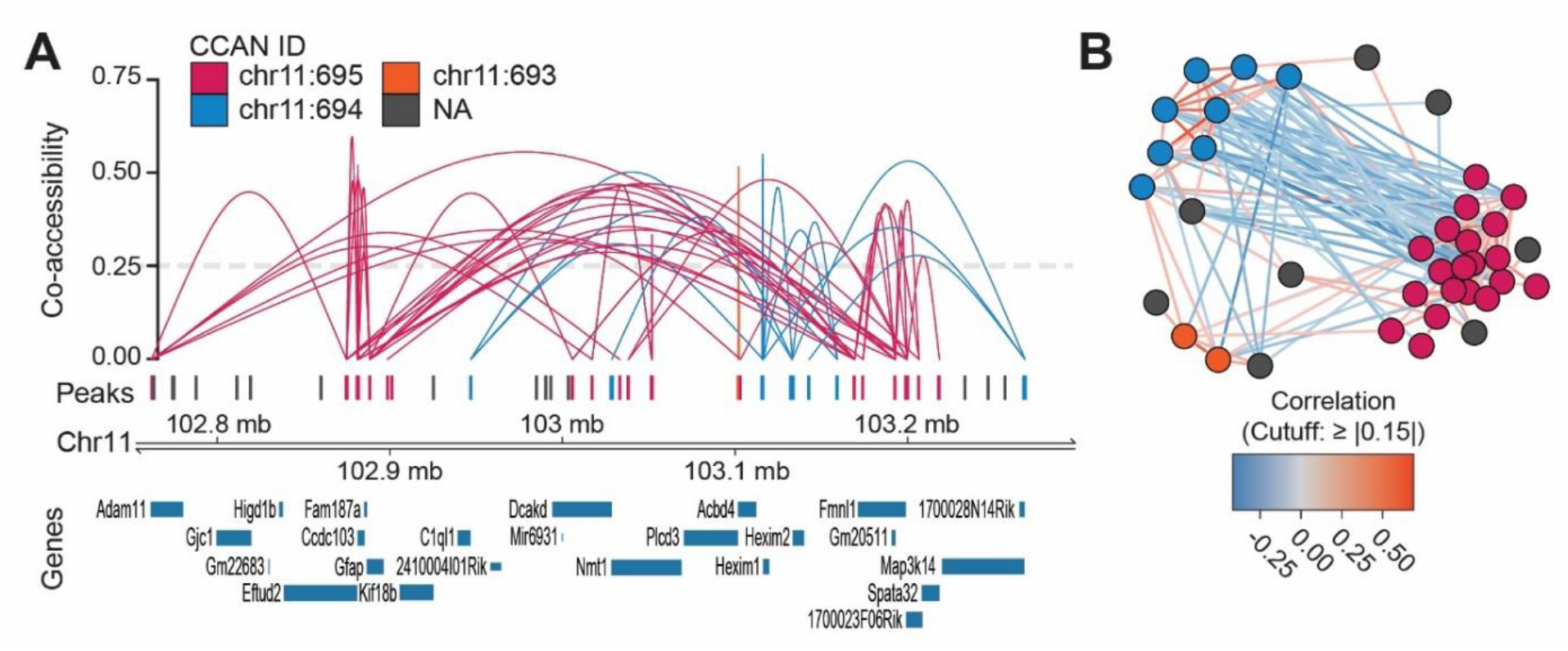
Mutually exclusive CCAN. A) Example of a mutually-exclusive set of CCANs that includes the astrocyte marker gene Gfap. CCAN 695 (pink) and 694 (blue) overlap one another in genome positions; however, they are comprised of mutually-exclusive peak sets, suggesting two alternative chromatin conformations. B) The mutually exclusive CCAN peak sets have negative correlations with peaks from one another’s network.

**Supplementary Figure 14:**
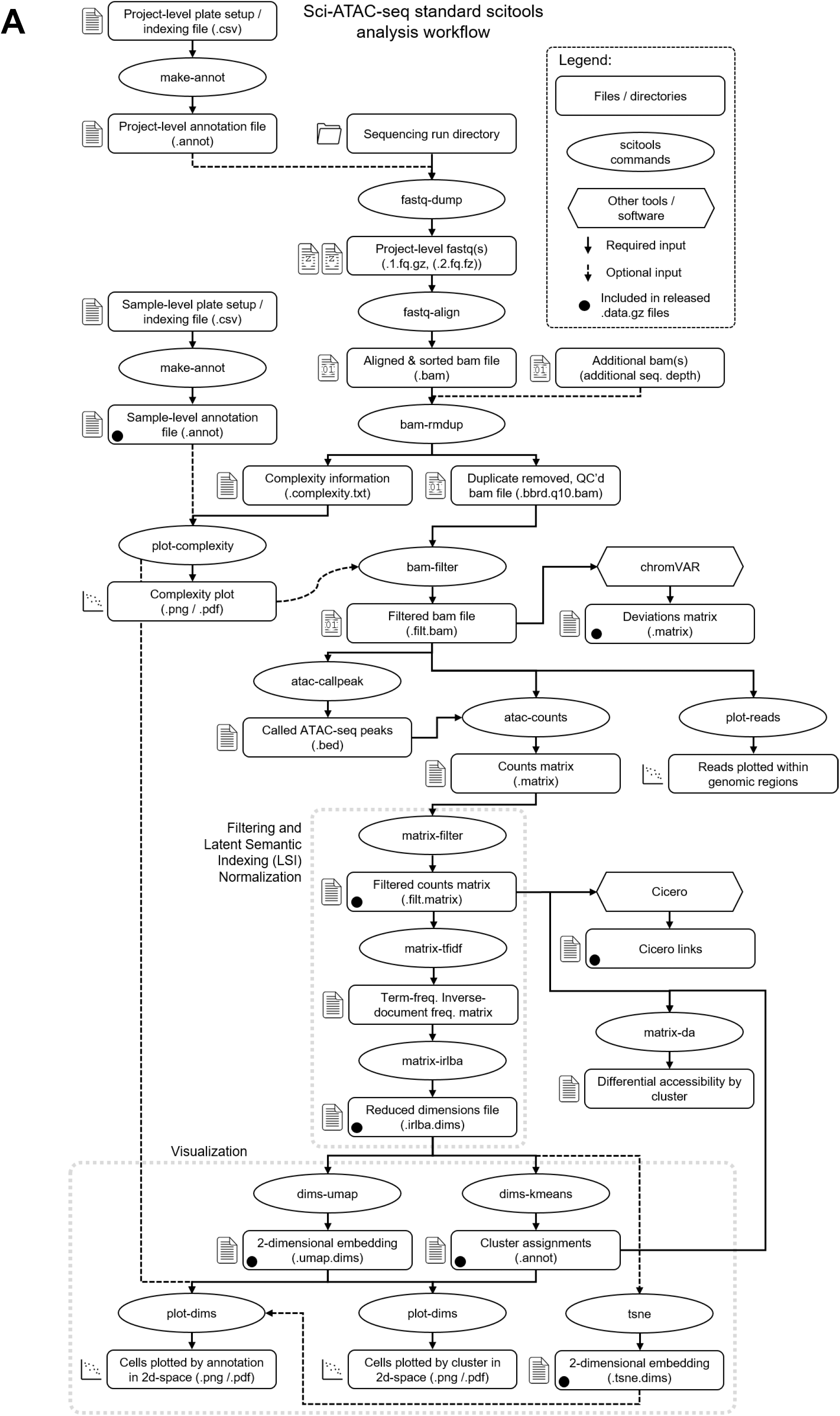
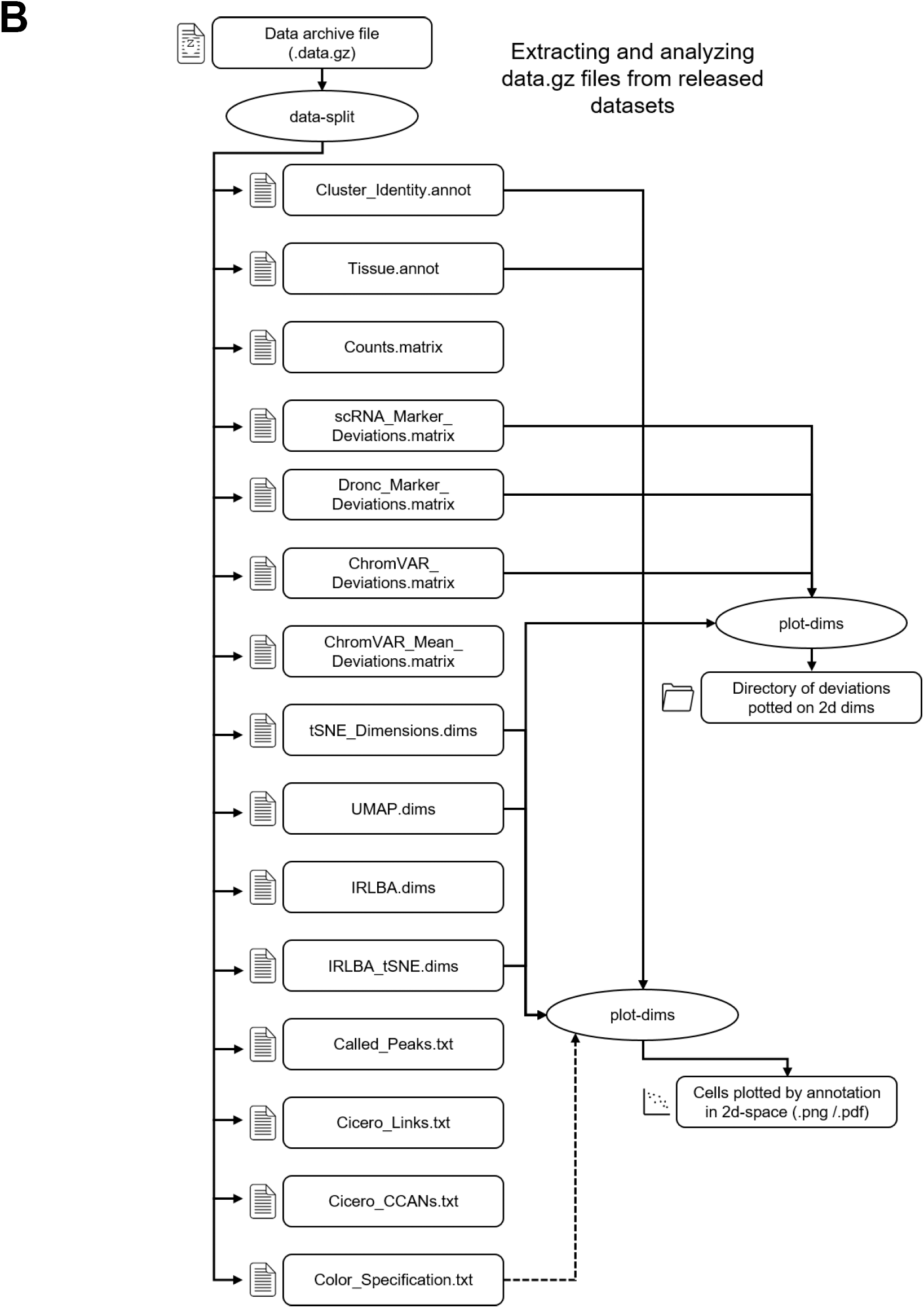
Scitools analysis and data access workflow. Detailed breakdown of all analyses performed by scitools in this study, and general best-practice guidelines for processing sci-ATAC-seq data using scitools. B) scitools data-split command to extract data files associated with this study from the GEO archive.

## Supplementary Note 1: Data & Software Description

We have released our data in a format compatible with the scitools software suite, which can be found at https://github.com/adeylab/scitools. For more information on scitools commands and usage, refer to user manual provided at the GitHub site.

To extract associated data, run the following command:

~~~
$ scitools data-split [dataset].data.gz
~~~

This will produce a set of files for each of the data sets with self-descriptive titles:

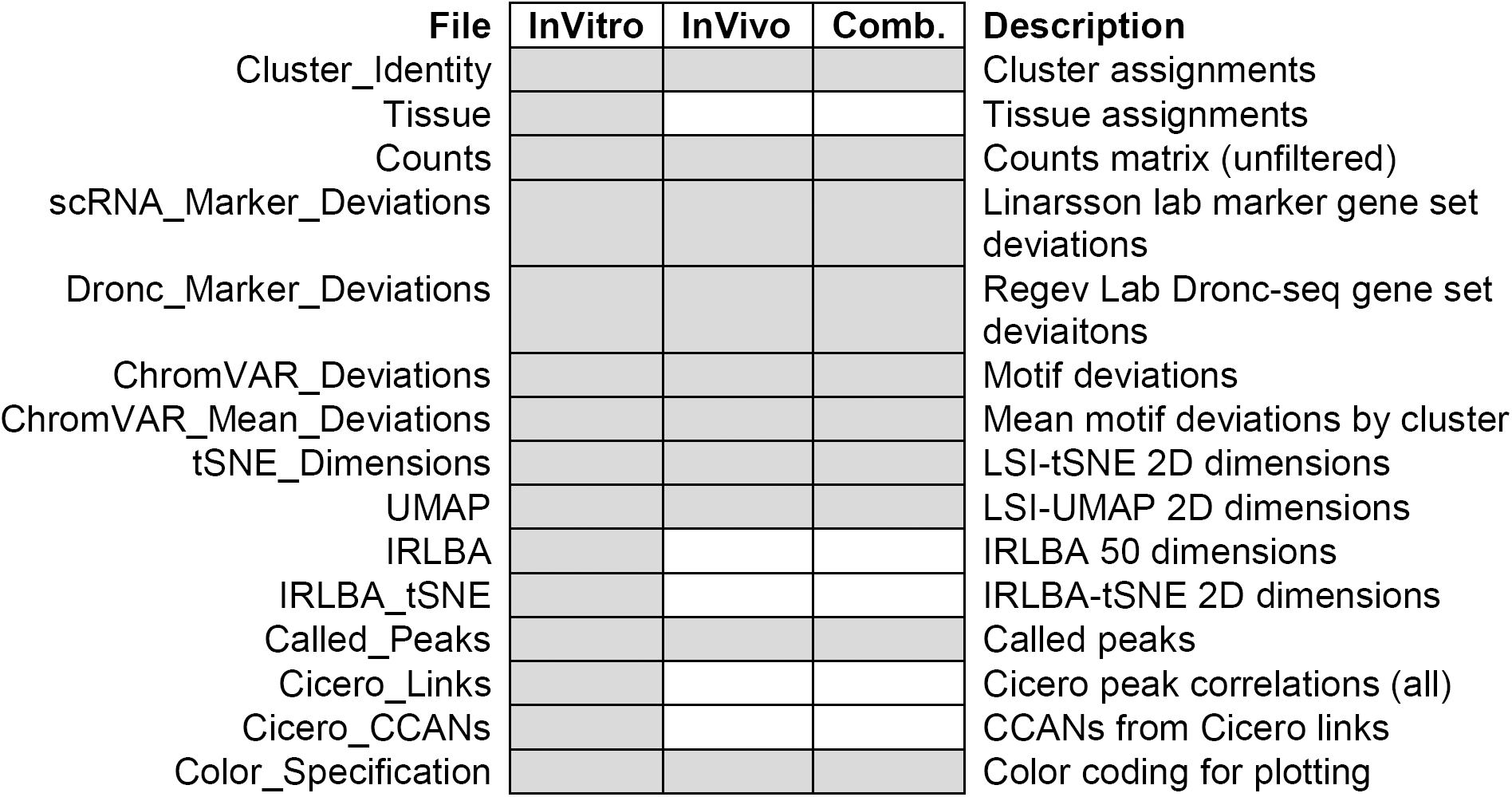

Scitools commands can then be run to reproduce figures in the manuscript. Below are several examples:

1. Plotting tSNE projections of cells:

~~~
$ scitools plot-dims –A [dataset].Cell_Type.annot [dataset].tSNE.dims
~~~
2. Plotting the chromVAR deviation scores onto tSNE dims:

~~~
$ scitools plot-dims –M [dataset].ChromVAR_DevZ.matrix [dataset].tSNE.dims
~~~

The counts matrix is also produced by the data-split command which can be used to run other scitools functions, for example:

1. Performing latent semantic indexing:
  a. Filter matrix:

~~~
$ scitools matrix-filter –C 1000 –R 50 [dataset].Counts.matrix
~~~
  b. Tfidf transform:

~~~
$ scitools tfidf [dataset].Counts.matrix
~~~
  c. Perform LSI:

~~~
$ scitools lsi [dataset].tfidf
~~~
  d. Perform tSNE:

~~~
$ scitools tsne [dataset].tfidf.LSI.matrix
~~~
2. Alternative dimensionality reduction strategies:
  a. On the tfidf matrix, perform irlba:

~~~
$ scitools irlba [dataset].tfidf
~~~
  b. Perform tSNE:

~~~
$ scitools tsne [dataset].tfidf.irlba.matrix
~~~

To list additional commands that can be run on the datasets, run the following:

~~~
$ scitools list
~~~

